# A standardized and reproducible behavioral protocol to elicit visual spatial attention in mice

**DOI:** 10.64898/2025.12.19.695467

**Authors:** Kayla Peelman, Nicole Allen, Jamin Ahn, Joseph Del Rosario, Kaitlin C. Jacobson, Esther Kim, Anthony D. Lien, Mercedes Lopez-Esteva, Lyndah Lovell, Kendell Worden, Yanlin Yang, James Zhuang, Bilal Haider

## Abstract

Understanding how neural activity gives rise to cognitive processes such as selective attention is a fundamental goal of neuroscience. An important but often overlooked advance towards this goal requires the development and sharing of rigorous and reproducible behavioral tasks across labs; this is particularly important given the recent surge in studies of cognition and perception in mice. Here, we developed a standardized training protocol for head-fixed mice to become experts in a psychometric visual contrast detection task in just 17 days. Experts detected stimuli at two distinct spatial locations for several hundred trials per day. As consecutive trials elapsed at either location, the speed, accuracy, and contrast sensitivity of visual perception improved – all hallmarks of spatial attention improving performance, as seen in primates. These findings validate the efficacy of this protocol to reveal multiple aspects of selective spatial attention in mice, establishing a rigorous and reproducible tool for the community.

## Introduction

One of the central goals of neuroscience is to determine the relationship between neural activity and behavior. This typically involves monitoring neural activity in animals while they perform a behavior of choice, then using analytical methods to relate the neural activity to behavioral features of interest. Such studies often report highly complex methods for neural activity measurements, data processing, data analysis, and statistical quantification – but with much less detail provided for the methodological decisions and procedures involved in the often equally complex behavioral tasks. When these details are provided, they often only describe end-stage, expert-level behavioral performance – not how animals were trained to achieve this state. This shortcoming in providing detailed behavioral training protocols alongside other reproducible deliverables of research (such as data and code reproducing the results) is a critical hindrance for the widespread replication of behavioral neuroscience studies. Indeed, recent large-scale, collaborative studies of visual perception have highlighted how standardizing and sharing behavioral protocols across labs improves reproducibility and interpretability of neural processes underlying behavior, a major step towards increasingly rigorous science^1^.

Many fundamental studies of neural activity underlying behavior have focused on visual perception, and how this process is affected by selective attention. Selective attention involves focusing processing to one aspect of the environment, while reducing processing of other aspects^2,3^. For example, selective attention to a particular region of visual space is accompanied by faster and more sensitive behavioral responses to stimuli in that region^4^. This selective improvement comes at the cost of slower responses and lower sensitivity to stimuli appearing elsewhere^5^. For many decades, the neural basis of this process has been studied in great detail in head-fixed monkeys performing selective attention tasks^6–10^. Only very recently was it shown that head-fixed mice can also use selective visual attention to improve their visual processing^11^. Although some visual specializations of primates are absent in mice, they nonetheless share many fundamental organizational principles of visual processing, including retinotopy and hierarchical, multi-area feedforward and feedback connectivity^12–16^. Several groups have now developed tasks where expert mice show many signatures of visual spatial selection and attention^11,17–23^. These and other tasks for mice hold great potential for the use of advanced technologies^24,25^ to study the neural basis of selective attention at scales that are more challenging to measure in primates.

Detailed reporting of behavioral training protocols for sophisticated selective attention tasks in mice is particularly important for interpretability of the underlying neural mechanisms. Mice can be idiosyncratic during learning and behavioral performance^26–29^, and sophisticated tasks can be learned and executed in different ways by experts, with distinct consequences for neural activity underlying performance^30^. It is challenging to draw firm conclusions about the generality of mechanisms underlying selective attention across subjects, across tasks, and across species, without standardized and reproducible training methods and rigorous performance tracking from naïve to expert stages. Thus, standardizing how mice are trained in selective attention tasks is critical for generating reproducible, interpretable inferences about this fundamental cognitive process.

Here we detail a standardized behavioral protocol for training head-fixed mice to become experts in a visual task that elicits several signatures of spatial attention. With an average of 17 days of training, mice could report detection of small, localized visual stimuli that appeared at varying contrasts in one of two locations in the visual field. The stimuli appeared in blocks of consecutive trials at one location, or the other. This alternating block structure elicited improvements in reaction time and accuracy as trials progressed at a given location, replicating our prior work^20^; additionally, our current protocol elicited clear measures of attentional improvements in both monocular and binocular visual space. Furthermore, mice trained with this protocol also improved their contrast sensitivity across successive trials at a given spatial location, just like attentional improvements seen in monkeys and humans. These new results further validate the efficacy of this protocol, establishing an efficient, high-throughput, and reproducible tool for the study of multiple aspects of selective spatial attention in mice.

## Results

### Behavioral task design and training phases

We previously developed a behavioral task for head-fixed, water-restricted mice that elicited both behavioral and neural signatures of visual spatial attention^20^. In that study, mice reached standardized expert-level performance, but without using a rigorously operationalized training protocol with standardized performance benchmarks during training. Here, we describe how standardizing and operationalizing key aspects of the training protocol into phases enables significantly faster learning to expert stages, and consequently more robust measurements of several signatures of visual spatial attention that are repeatable across days and across subjects. The optimized protocol replicates our prior work, and now improves training efficiency to increase throughput for the study of visual attention in mice.

The end goal of the task is to measure how expert mice use spatial attention to improve the speed and accuracy of visual perception; we will first briefly describe the expert stage of the task, to contextualize the subsequent descriptions of training stages. Expert mice report their perception of small, static visual gratings (shown at a range of contrasts; see Methods) by licking for water rewards. Rewards are only available during a brief response window (1 s) while the stimuli are on the screen (Fig. 1A); the first lick during the response window triggers reward delivery. Some trials are 0% contrast (blank) to measure the false alarm rate. Each detection trial is preceded by a mandatory quiescence period of randomized duration (0.5 - 15 seconds of blank screen; Fig. 1A), where stray licks restart the trial; this incentivizes mice to withhold licking until stimulus appearance, then to lick rapidly upon stimulus detection. The key feature of expert-level task performance is detection of stimuli that appear in different spatial locations across blocks of consecutive trials. For example, in one block of consecutive trials, the stimuli appear in front of the mouse (Fig. 1B, binocular block, blue). Then, without warning, the stimuli appear in a peripheral spatial location for the next block of consecutive trials (Fig. 1B, monocular block, magenta). Aligning behavioral performance to the first trial of the block (when the stimulus switches from the expected to unexpected location) allows us to examine how perceptual speed, accuracy, and sensitivity change across consecutive trials in the block. If the block structure elicits spatial attention, we should observe that perception is hindered on the first trial in the block (when attention is elsewhere), but then improves across consecutive trials at the new location.

**Figure 1.**
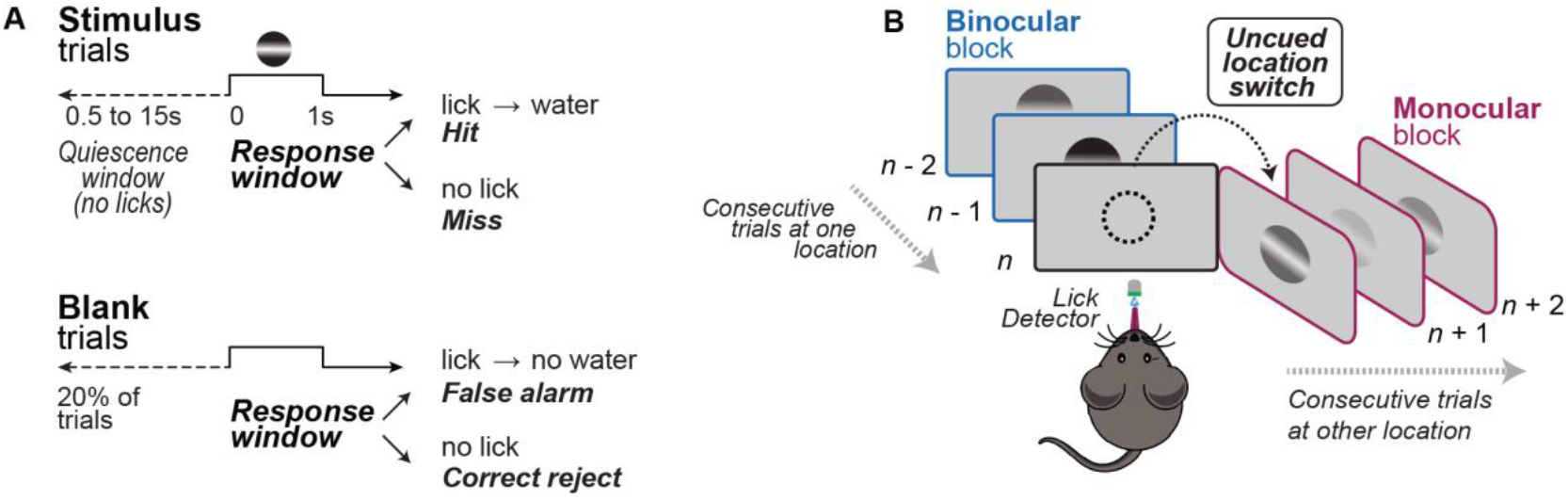
Training head-fixed mice in phases to elicit the behavioral correlates of visual spatial attention. **A**. Head-fixed, water-restricted mice reported stimulus detection by licking for water. All trials started with a quiescence period of randomized duration (0.5 - 15 s). Licking during this period restarted the trial. After stimulus onset, the first lick during a 1 second response window dispensed water (hit trial); failure to respond resulted in no reward (miss trial). On 20% of trials (bottom, blank trials), a 0% contrast stimulus was presented. Licking during these trials was recorded as a false alarm, while withholding licking was considered a correct rejection. **B**. At the expert stage, stimuli appeared in blocks for 10-30 consecutive trials at a single spatial location (binocular or monocular visual field). At the end of a block of trials in one location, the stimulus switched to the opposite spatial location without a cue.

Overall, averaged across all subjects trained with this protocol (N=36), we observed clear behavioral effects of spatial attention (Fig. S1A). Both detection speed and perceptual sensitivity improve significantly across consecutive trials of stimulus detection in peripheral (monocular) space, consistent with our previous study^20^. However, our prior study did not probe the effects of attention in the binocular visual field; here, we show the training protocol also elicits clear effects of visual attention in the binocular visual field (Fig S1B), as will be detailed later.

### Training protocol with phases enables faster task expertise

The training protocol consisted of five distinct phases (Fig. 2A), each designed to systematically develop task proficiency and ultimately reveal behavioral signatures of visual spatial attention. We briefly describe the phases here, then in greater detail in subsequent sections.

**Figure 2.**
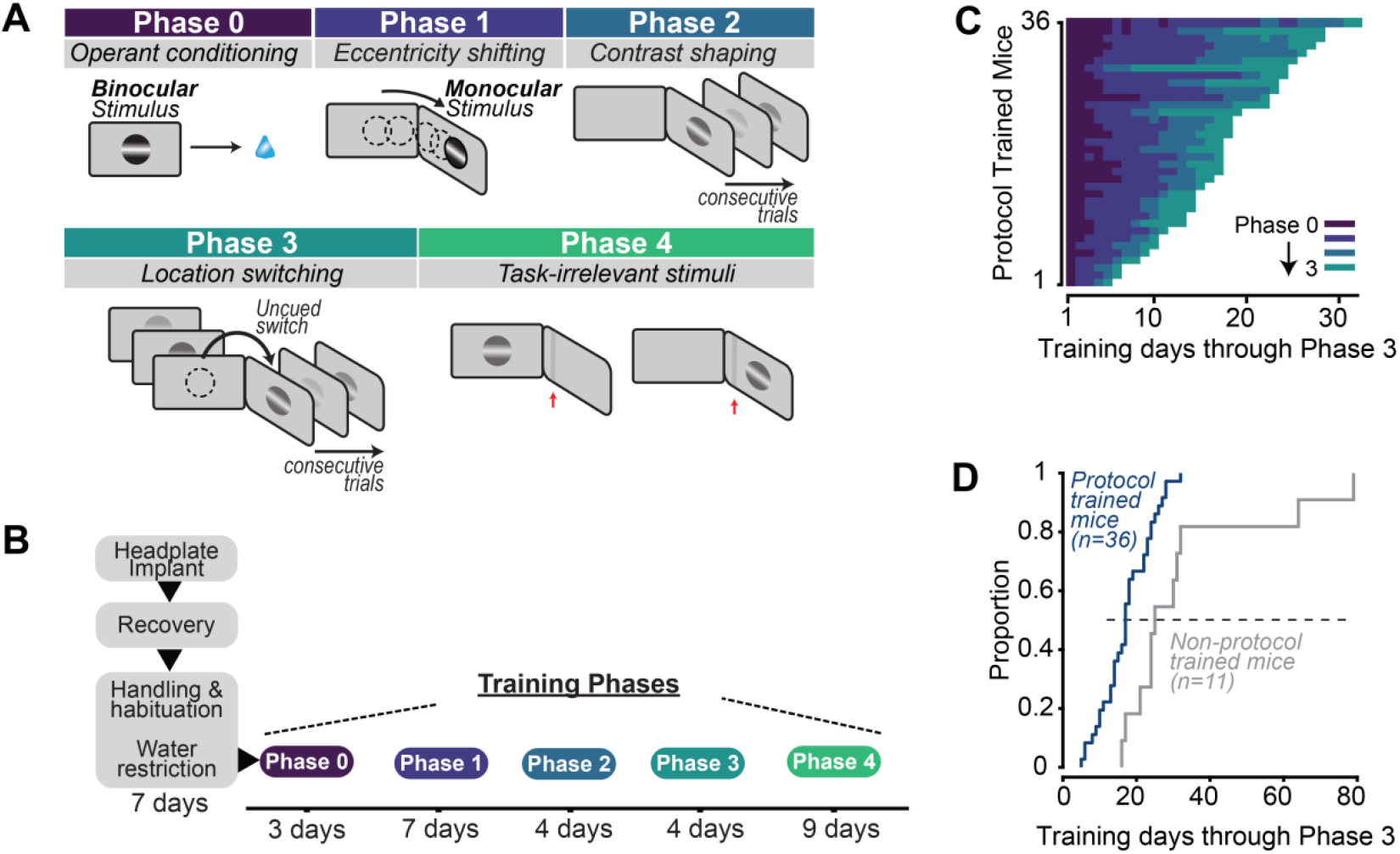
Mice trained with a phased protocol become experts faster. **A**. Schematic of sequential training phases: Phase 0 (conditioning); Phase 1 (shifting stimulus eccentricity); Phase 2 (contrast shaping); Phase 3 (switching stimulus locations); and optional Phase 4 (introduce task-irrelevant stimuli). Attentional effects appear in expert mice from Phase 3. **B**. Timeline of training phases, with average days per phase across mice. After surgical implantation, recovery, handling and habituation plus water restriction (7 days total; last 1-2 days introduce water restriction), protocol trained mice took another 17 ± 7 days (mean ± SD) to become experts in the task (N = 36 mice). Phase 0 (3 ± 2 days, mean ± SD); Phase 1 (7 ± 4 days); Phase 2 (4 ± 3 days); Phase 3 (3 ± 3 days); optional Phase 4 (9 ± 11 days). **C**. Number of days each mouse spent in phases 0-3 (color), rank ordered by total training duration. **D**. Mice trained in protocol (dark blue) become experts in 17 ± 7 days, twice as quickly as mice trained without a standardized, phased protocol (gray; 33 ± 20 days; *p* = 0.017, Kolmogorov-Smirnov test).

In Phase 0 (operant conditioning), water-restricted mice were introduced to the task by associating high-contrast stimulus appearance with automatic delivery of water rewards (0.7 seconds after stimulus onset); this positively reinforces licking following visual stimulus onset. The stimulus is only shown in one location in Phase 0 (at the vertical meridian in the binocular zone, defined as 0°) until mice show reaction times (first lick latencies) that are tightly aligned after visual stimulus onset and before the automatic reward delivery (anticipatory licking); thereafter, the automatic reward is discontinued and the first lick during the visual stimulus triggers reward delivery.

Phase 1 (eccentricity shifting) established stimulus detection at progressively greater distances from the center of the visual field. Eccentricity refers to the angular distance of a visual stimulus relative to the center of gaze, typically in horizontal space. In mice, stimuli with eccentricity >40° away from the vertical meridian (defined as 0°) appear in the monocular visual field. We gradually moved the stimulus location in 10-20° increments towards the monocular field (up to 70 - 90° eccentricity), with progression determined by performance criteria (hit rate ≥75%, d prime > 1). In Phase 2 (contrast shaping), stimuli remained fixed at 70°-90° but were shown at varying contrasts. This ensures mice learn to detect stimuli of different intensities and allows assessment of psychometric performance spanning the detection threshold.

Phase 3 (location switching) introduces contrast-varying stimuli in two different locations (either 0° or the peripheral location), in blocks of 10-30 consecutive trials at a given spatial location. During each behavioral session, trial blocks switch from one location to the other. Phase 3 allows concurrent assessment of both psychometric performance and spatial attention effects elicited by block switching.

Finally, in mice eventually used for neural recordings, we introduced an optional Phase 4 (Task-irrelevant stimuli). Mice continue performing the task exactly as described in Phase 3, but in the presence of low contrast, task-irrelevant vertical bars flashed across the visual field. These task-irrelevant stimuli are used to assess attention-related changes to neural spatial receptive fields^20^. This stage is not required to assess the behavioral effects of spatial attention induced by location switching.

We hypothesized that this structured, phased approach would expedite task learning compared to our prior training methods. Indeed, 92% of mice designated for training with the phased protocol (36/39; see Methods for inclusion criteria) became experts in just 17 ± 7 days (Fig. 2B, C; through Phase 3). This was a 49% reduction in training time compared to our previous methods (Fig. 2D; *p =* 0.017, Kolmogorov-Smirnov test). Mice trained with the phased protocol also showed significantly less variability in total training time relative to the non-protocol trained mice (N=11 mice, *p =* 0.033, Brown-Forsythe test). Expert mice trained with the protocol performed 21 ± 6 spatial location switches per day, and performed 369 ± 78 trials (mean ± SD), providing hundreds of thousands of standardized behavioral trials in a large cohort of mice for detailed analysis of attentional effects, as will be shown later. In the next sections, we detail the quantitative criteria used for progression through each of the training phases.

### Operant conditioning establishes stimulus-reward association

Mice learned the association between visual stimuli and rewards through operant conditioning (Phase 0, Fig. S2). During this phase, after a randomized blank screen period (1 – 15s, see Methods), static horizontal Gabor gratings (65-85% contrast) appear in the binocular visual field (0°, azimuth) for 2 seconds. Water rewards were delivered automatically 0.7 seconds after stimulus onset to encourage mice to associate the visual stimulus with water delivery. Mice showed evidence of learning the stimulus-reward association with a progressive shift of first lick latencies from times after reward delivery (Fig. S2A, B; 0.93 ± 0.05 s, first day of Phase 0 training) to times before reward delivery (Fig. S2B; 0.55 ± 0.08 s, N=36 mice, median ± IQR/2, final day of Phase 0 training, *p =* 2.56e-7). They concurrently showed a higher probability of trials with lick responses across the 3-9 days of Phase 0 training (Fig. S2C, D; from 52 ± 10% of trials to 97 ± 8% of trials on first to final day of Phase 0 training; N=36 mice, median ± IQR/2). This shift in the timing and probability of licking indicates a transition from reward-driven licking to visual-driven licking. On the last day of Phase 0, automatic reward delivery was discontinued, and reward was only delivered if a lick occurred during visual stimulus presentation, thus directly reinforcing the visually elicited action. Mice advanced to Phase 1 when these lick times occurred consistently before 0.7 s with low trial-to-trial variability (Fig. S2 C-F). Thus, at the end of Phase 0, licks report perception of the visual stimulus itself.

### Stimulus detection across the visual field and at varying contrasts

In Phase 1, we shortened the response window (to 1 s) to encourage fast reaction times, and gradually moved the stimulus from the central to peripheral visual field (Fig. 3A). As shown in our previous study and several others^31–33^ mice show greatest visual sensitivity in the binocular visual field (spanning ± 40 degrees of central visual space) (Fig. S3A, B). Accordingly, the stimulus becomes more difficult to detect as it shifts in eccentricity away from the vertical meridian. This necessitates gradual, stepwise introduction of the stimuli in more challenging monocular locations, with concurrent assessment of performance as a function of stimulus position. The stimulus is shown at a fixed position in a block of trials (typically 50 – 100), and then shifted by 10 – 20°, pending performance metrics (hit rate > 75%, d’ > 1). It is important to enforce low false alarm rates in Phase 2 (captured by d’) before moving the stimulus position. In an example mouse (Fig. 3B), stimuli placed in the binocular zone resulted in high detection accuracy (HR > 80%). However, as the stimulus shifted in eccentricity across sessions, accuracy often abruptly decreased; further, if stimulus eccentricity shifted too suddenly (i.e., before achieving high performance at a given location), mice often disengaged from the task or would not perform consistently at more eccentric stimulus positions (Fig S4). However, with repeated exposure at these locations, performance stabilized and improved. A similar trend was observed across the population (Fig. 3C): performance remained consistently high using gradual, stepwise shifts in stimulus eccentricity. Importantly, these improvements were not accompanied by significant increases in the false alarm rate (Fig S3C). These results highlight the importance of gradually shifting the eccentricity of the stimulus for ensuring high detection performance at these more difficult spatial locations.

**Figure 3.**
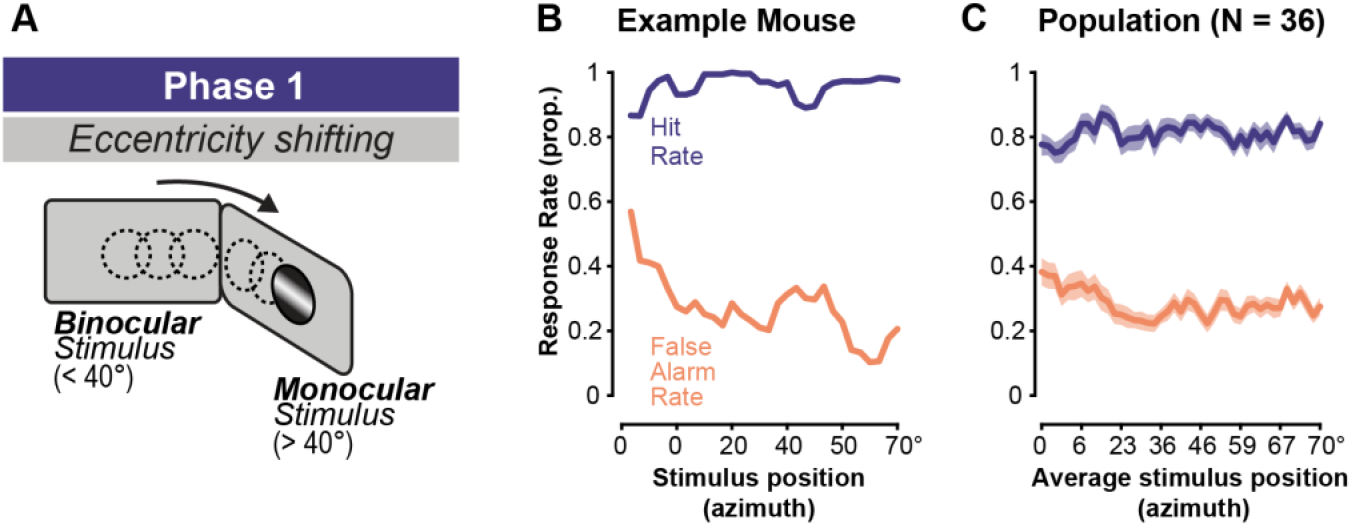
Shifting stimulus eccentricity from central to peripheral visual field. **A**. Grating stimulus is presented at a fixed location for a block of trials. Across blocks of trials, grating location is slowly moved in increments of 10-20° towards the monocular field of view. **B**. Performance in an example mouse. Slow progression of stimulus from the binocular to monocular visual field keeps performance high. False alarm rate continues to decrease while hit rate remains stable. **C**. Population of mice trained in phases (N = 36). Data plotted are mean ± SEM across mice. On the final day of Phase 3 where the stimulus is at 70°, the hit rate is 82±12% (mean ± SD) and false alarm rate is 27±11%.

Phase 2 training introduced stimuli of different intensities and measured psychometric detection sensitivity (Fig. 4A). After establishing high detection performance for high contrast stimuli in the far periphery (monocular space; ~70°), we next systematically varied the range of contrast levels across sessions while keeping the stimulus fixed at this location (4-6 different contrasts per session; 100-150 trials per session). This has two purposes: 1) it prevents behavioral ceiling effects (i.e., an inability to measure attentional improvements beyond maximal performance at high contrast), and 2) it ensures that mice are performing around their detection thresholds, creating conditions that facilitate task engagement so that attention may improve the speed and probability of attaining rewards. During the early sessions of Phase 2 (first 25% of sessions), a range of medium to high contrasts appeared (e.g., 18, 33, 65, 85%), resulting in high performance (>80% correct) across all stimuli (Fig. 4B, orange). In the subsequent sessions, when performance at the highest contrasts exceeds 80% or when responses at the lowest contrast rise significantly above the false alarm rate, the contrast range is shifted downward by one contrast step, ensuring that responses sample the full dynamic range. As a result, the lowest contrasts in early sessions become the highest contrasts in the later sessions. In late sessions of Phase 2 (last 25% of sessions), this adaptive approach reveals psychometric performance that spans from the false alarm rate to maximal hit rates (Fig. 4B, blue circles). In addition, late Phase 2 is characterized by improved performance at intermediate and low contrasts (Figure 4B, C). Population analysis revealed a significant reduction in contrast detection threshold in late versus early Phase 2 training (Figure 4D; early: 18.6 ± 9.6 %; late: 5.8 ± 2.3%; *p =* 4.9e-5; median ± IQR/2, sign test).

**Figure 4.**
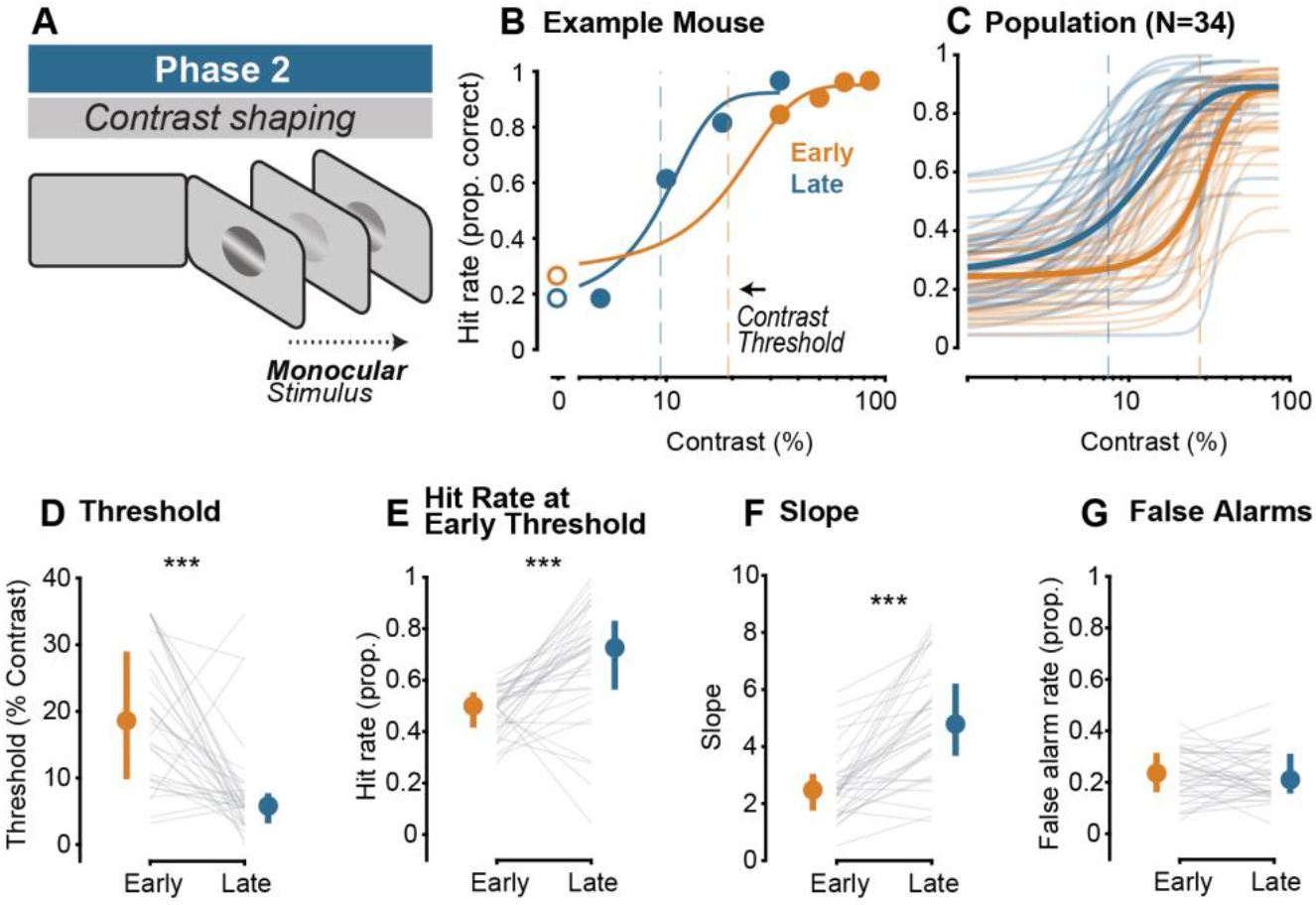
Shaping improves psychometric contrast detection at monocular locations. **A**. Schematic of Phase 2: Contrast shaping. Grating stimulus stays in the monocular location (70°) but appears across a range of contrasts (4-6 different contrasts, not including 0%). **B**. Early (orange) vs Late (blue) psychometric performance during Phase 2 of training for one example animal. Note that in early contrast shaping, performance is high for all contrasts, while in late contrast shaping, the mouse exhibits psychometric performance at a range of contrasts that evenly sample the animal’s psychometric range (N=1 mouse; early R^2^: 1.00; late R^2^: 0.90). Dashed lines indicate contrast threshold for Early (orange) and Late (blue) curves. Curves fit using a Boltzmann equation (see Methods) throughout figure. **C**. Early (orange) vs late (blue) psychometric performance during Phase 2 of training for entire population (N=34; early average population R^2^: 0.91 ± 0.21; late R^2^: 0.91 ± 0.17). Note that two animals were omitted because of only 1 day of Phase 2 training, preventing ‘Early’ and ‘Late’ comparisons. **D**. Contrast threshold decreases from early to late Phase 2 training (early: 18.6 ± 9.6 %; late: 5.8 ± 2.3%; *p =* 4.9e-5, median ± IQR/2, sign test throughout figure unless otherwise noted). Each gray line indicates an individual animal’s change in contrast threshold from early to late Phase 2. Error bars indicate median ± IQR throughout figure. **E**. Hit rate improves at lower contrasts across training in Phase 2. Threshold contrast was established during early training (50.0% ± 6.8% accuracy) and hit rate at that same contrast was calculated during Late Phase 2 training (72.6 ± 13.3% accuracy at same contrast value; *p =* 1.1e-4). **F**. The local slope of the psychometric curves significantly increases from Early (2.48 ± 0.64) to Late (4.79 ± 1.26) sessions in Phase 2 training (*p =* 1.15e-6). **G**. False alarm rate (FAR) does not significantly increase across phase 2 training (early: 23.3 ± 8% FAR; late 20.7 ± 8% FAR; *p =* 0.845).

Hit rates at a “reference” intermediate contrast value (defined as the threshold contrast in Early sessions) rose from 50.0% ± 6.8% to 72.6 ± 13.3% (Fig. 4E; *p =* 1.1e-5, sign test). Similarly, the slope of the psychometric curve becomes steeper from early to late Phase 2 (Fig. 4F, early slope: 2.5 ± 0.65; late slope: 4.8 ± 1.3; *p =* 1.15e-6; median ± IQR/2, sign test). This steeper slope indicates improved sensitivity, such that smaller changes in stimulus contrast around threshold produce larger changes in performance. At the same time, false alarm rates did not change across Phase 2 training (Fig. 4G; early: 23.3 ± 8%; late 20.7 ± 8%; *p =* 0.845). These perceptual improvements validate the effectiveness of our structured, multi-phase training approach for optimizing psychometric visual detection. Once mice reached this stage of high-quality psychometric performance in peripheral stimulus space, we introduced stimulus location switching to assess spatial attentional effects.

### Stimulus location switching elicits spatial attentional effects on behavior

Stimulus location switching (Phase 3) provoked behavioral effects consistent with spatial attention, in both binocular and monocular visual fields. We first ensured psychometric performance throughout the visual field during location switching. Unlike monocular detection, which requires gradual shaping for high psychometric performance, binocular detection is both intrinsically easier and more contrast sensitive (Fig. S3), so we can immediately re-introduce binocular stimuli at different contrasts, but at a range one step lower than the monocular stimuli. This lower range spans the full dynamic range near chance level detection to maximal performance (Fig. 5A). During location switching, we measured high quality psychometric performance in both locations for all mice trained with the phased protocol (Fig. 5A; binocular psychometric curve fit quality R^2^ = 0.97 ± 0.05; monocular R^2^ = 0.95 ± 0.12, mean ± SD, N = 36 mice). This defines expert-level task performance.

**Figure 5.**
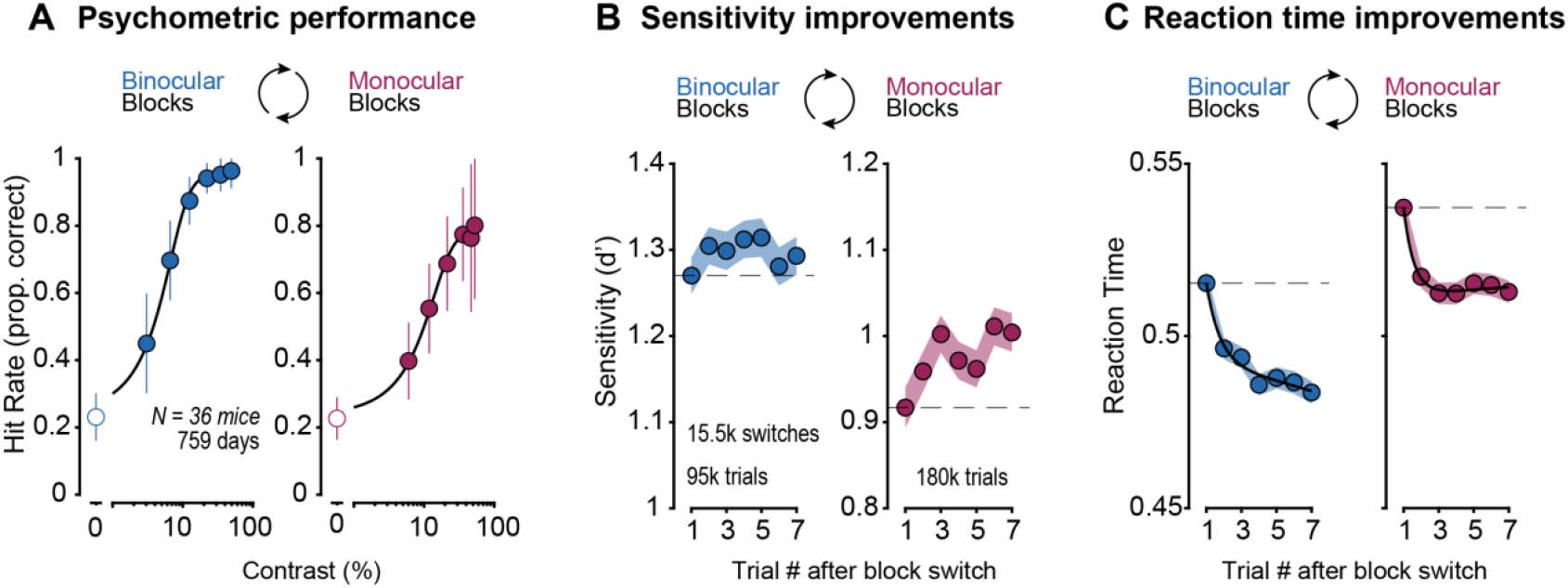
Stimulus location switching reveals behavioral effects consistent with spatial attention in both binocular and monocular spatial locations. **A**. Psychometric performance is observed in all mice (N = 36) for both binocular (left, blue) and monocular (right, magenta) spatial locations (Boltzmann fits; binocular R^2^ = 0.99; monocular R^2^ = 0.99; figure error bars represent mean ± SD across N = 36 mice, 759 days, 15,617 total block switches, 95,295 total binocular trials, 180,617 total monocular trials; on average mice executed 21 ± 6 block switches, and performed 369 ± 78 trials per day. Each mouse contributed 21 ± 11 days of data; mean ± SD). All eligible days in Phases 3 and 4 were analyzed (no sub-selection). **B**. Sensitivity improves significantly as a function of trial number in block for monocular stimuli (right, magenta; *p =* 6.0e-5, linear mixed effects model throughout figure, see Supplementary Table 1). Binocular sensitivity improves after Trial 1, but not significantly (left, blue; *p =* 0.64). Data are plotted as mean ± SEM across days. Note different scales for ordinate. **C**. Reaction times become significantly faster as a function of trial number in block for both binocular (left, blue; *p =* 2.9e-30) and monocular stimuli (right, magenta; *p =* 2.9e-11). Data are plotted as mean ± SEM across days.

As mice detected stimuli spanning psychometric thresholds, switching the stimulus location across blocks elicited attentional improvements of overall detection sensitivity (d’) across successive trials (Fig. 5B). Sensitivity improved after the first trial in either location, but significantly for monocular stimuli (*p=* 6e-5, linear mixed-effects model, n = 5313 observations from Phase 3 and Phase 4 in 36 mice; see Methods and Supplementary Table 1 for model details).

Simultaneously, reaction times significantly decreased across these same trials in both locations (Figure 5C, monocular: *p =* 2.9e-11; binocular: *p =* 2.9e-30;). Restricting the analysis to Phase 3 alone (20% of Fig. 5 dataset) showed that the effects were already present from the start of location switching (monocular d’: *p =* 0.001, binocular d’: *p =* 0.285; binocular RT: *p =* 1.86e-08; monocular RT: *p =* 0.003; linear mixed effects model). Increased sensitivity and faster reaction times across trials were evident in the majority of mice trained in the protocol: 83% exhibited at least one day with significant behavioral effects of attention during Phases 3 and 4, and 61% of individual subjects showed significant effects in the average of all their Phase 3 and Phase 4 sessions (no selection criteria; Fig. S5). Importantly, these behavioral improvements were not associated with changes in eye position or pupil area across trials following block switches (Fig. S6), and without increased false alarms during location switching (Phase 2 = 23% ± 10%; Phase 3 = 23% ± 6%; Phase 4 = 24% ± 6%; mean ± SD; *p =* 0.350, Kruskal-Wallis test). Together, these findings confirm that just like prior versions of this same task^20^, location switching across blocks reliably elicits spatial attentional effects; additionally, the new phased protocol elicits attentional effects in both spatial locations, all while mice detect stimuli at a range of contrasts throughout the visual field; this new feature of the training protocol allowed us to rigorously assess the potential for attentional effects on psychometric contrast sensitivity, examined next.

### Standardized training protocol reveals improved contrast sensitivity with attention

Having established repeatable effects of attention on detection speed and accuracy across subjects, we wondered if this cohort of expert mice could reveal additional effects of attention that were not explicitly designed into the protocol. Inspired by observations in monkeys and humans^34,35^, we assessed if attentional improvements in detection speed and accuracy were accompanied by heightened sensitivity to contrast. To test this, we aggregated and aligned all the trials following block switches in our protocol trained mice, selecting days with overall increasing d’ across successive trials following block switches (monocular: N=36 mice, 234 days, 808 block switches; binocular: N=36 mice, 254 days, 834 block switches). We then generated psychometric curves as a function of trials elapsed after the block switch. Recall that contrasts on every trial were randomized — trial 1 could be 10% contrast in one block, 18% contrast in another, etc. Thus, aggregated across all blocks and mice, the first trials of the block switches collected responses from all contrasts. The responses for all these 1^st^ trials following the block switch (spanning all contrasts) were sorted to construct a population level psychometric curve (Fig. 6A, D, black); the same procedure was done for the subsequent trials 2 - 5 (Fig. 6A, D). These block-switch aligned psychometric curves were constructed separately for each spatial location, showing proportion correct as a function of contrast and as a function of trials elapsed across the block.

**Figure 6.**
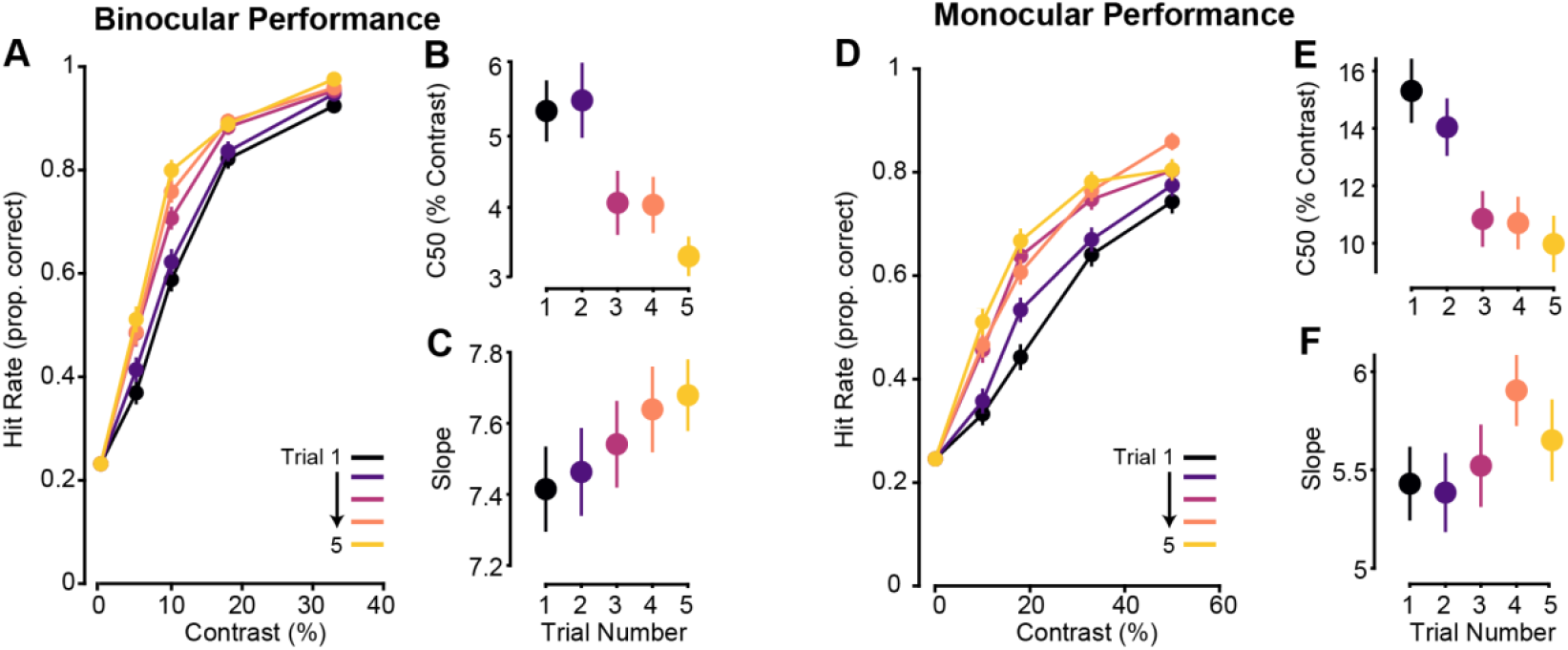
Contrast sensitivity improves after stimulus appears in new location. **A**. Psychometric curves for binocular detection showing hit rate as a function of stimulus contrast for trials following a block switch. Data from days with overall increasing d’ across successive trials following block switches in phases 3 and 4 were aggregated for each trial number (1-5) after the switch into the binocular condition. Data are plotted as mean ± SEM across days throughout figure. **B**. Contrast threshold (C50), derived from trial-specific psychometric curves, was measured for binocular blocks (see Methods for defining contrast threshold). Following a switch to the binocular location, contrast threshold decreased significantly as a function of trial number in the block (linear mixed-effects model throughout figure, see Supplementary Table 2, *p =* 1.17e–6.). **C**. The slope of the contrast response function increased significantly as a function of trial number within block (*p =* 0.032). **D**. Psychometric curves as in A, but when the stimulus switched to the monocular location. **E**. Following a switch to the monocular location, contrast threshold significantly decreased as a function of trial number in the block (*p =* 2.38e–6). **F**. The slope of the contrast response function increased but not significantly as a function of trial number within block (*p =* 0.16).

This analysis revealed that, indeed, spatial attention also improved contrast sensitivity across successive trials in the block. Contrast thresholds and slopes of the psychometric function at threshold were analyzed as in a prior study^34^. Contrast thresholds significantly decreased across trials in both spatial locations (Fig. 6B, E; linear mixed-effects model: binocular: *p =* 1.2e-6; monocular: *p =* 2.3e-6). Similarly, the slope of each trial-derived psychometric curve increased as a function of trial number within a block, significantly for binocular stimuli (Fig. 6C, F; binocular: *p =* 0.032; monocular: *p =* 0.158). These results establish that contrast sensitivity in mice also improves in a manner consistent with effects of spatial attention in primates. These novel findings appeared in a large cohort of mice trained in a standardized protocol that was optimized to reveal effects of attention on two other behavioral correlates — improved reaction times and accuracy. These new results on improved contrast sensitivity further validate our protocol’s efficacy for instantiating multiple attentional effects on visual perception in mice.

## Discussion

Here we present behavioral data from a standardized protocol in which head-fixed mice become experts in a challenging psychometric visual task that elicits several signatures of spatial attention. Our protocol was explicitly designed to reveal changes in visual detection accuracy and reaction time as readouts of attentional improvements. We measured how these two quantities changed following the appearance of the task-relevant visual stimulus in an unexpected spatial location. Both quantities improved as consecutive detection trials elapsed at the newly relevant spatial location, consistent with well-known effects of spatial attention in monkeys and humans^2,3,5,7,34,36–38^. Our protocol showed significant improvements from our prior training procedures: task acquisition, expert-level performance, and attentional effects were repeatable across mice, and these were achieved in significantly fewer training days. Furthermore, by rigorously enforcing psychometric contrast detection in two spatial locations, expert mice trained with the protocol also showed improved visual contrast sensitivity across successive trials at a given spatial location, consistent with findings in monkeys and humans. Taken together, these findings establish and validate an efficient, high-throughput, and reproducible behavioral protocol for the study of spatial attention in mice, adding a valuable community resource for replicable behavioral neuroscience research.

The first advance of our protocol is that training times were fast and reliable. Mice needed just 17 days of training on average to achieve expert-level performance (defined as high-quality psychometric detection during location switching), cutting in half the required training days from our prior work. This time to expert-level performance is faster than other selective visual attention tasks in mice that report training times, ranging from 5 – 8 weeks^21^ up to 5 – 7 months^17,39^. Further, the variability of training days to reach expert stage was markedly reduced versus our prior work with a less operationalized training regime for the same task. Predictability of training durations is particularly important for experiments with time-sensitive variables (such as opsin expression, developmental milestones, studies of aging, chronic implantations, or studies with mouse models of neurological disorders that show age-related symptoms).

The second advance of our protocol is high efficiency of task acquisition and expert-level performance. Only 3% of mice (1/39) failed to reach expert level due to poor behavioral performance. Expert mice performed nearly 400 trials per day, and switched the location of visual detection 21 times per day. Several recent studies of visual attention in mice provide some comparison points for our protocol efficacy. In one study of audio-visual spatial attention, 37% (of 51 subjects) reached expert-level performance^17^. In another study, around 18% of mice were removed from training for failure to learn the task within 3 months^11^, but the remaining mice show high incidence of attentional effects within subjects. Another study of visual attention in freely moving mice showed between 11 – 32% exclusion of subjects due to poor task performance^19^. Our protocol efficacy compares favorably with these prior reports. However, it is important to note that we purposefully defined expert-level performance solely from psychometric criteria during location switching – not by the presence (or absence) of attention effects. This was to avoid experimenter bias in shaping the very effects we set out to test. Expert-level performance could certainly be achieved and maintained without requiring attention, since our task did not overtly cue or direct it. Nevertheless, location switching elicited multiple behavioral metrics that are consistent with effects of spatial attention improving performance.

Our gradually phased protocol with progressive increases of difficulty and session-by-session monitoring helped mitigate biases wherein mice favor trials in the binocular visual field, since this region is inherently much more perceptually sensitive^21,22,31–33^. Recent studies of visual selection and attention show that mice can adopt idiosyncratic biases for visual field preferences^18,21^, necessitating subject exclusion/grouping to average across mice with similar biases. Our protocol revealed clear population-level evidence for behavioral improvements in detection speed and contrast sensitivity in both the monocular and binocular visual fields, while detection accuracy improved in both locations but significantly in the monocular visual field. Importantly, these population-level findings on attentional improvements of accuracy, reaction times, and contrast thresholds were evident from the same interleaved behavioral sessions (with sequential location switching across both locations), and without sub-selecting only those subjects with statistically significant behavioral effects, or only selecting trials from one location or the other. Prior studies of selective spatial attention in mice that briefly cued the relevant visual hemifield^11^ established that this improves psychometric sensitivity to ensuing stimulus orientation changes. To our knowledge, our findings are the first to show in mice that spatial attention in either the central or peripheral visual field is associated with improved contrast sensitivity simultaneously with faster and more accurate detection.

Our training protocol offers several advantages for ease of implementation and scalability. First, mice were motivated only through water restriction, reducing complications associated with food restriction (preparation, delivery, and cleaning of liquid food; digestive complications, etc.). Water delivery is easy to implement, calibrate, and measure in real time, and provides an easy way to equalize motivation across subjects (body weight criteria provide an excellent proxy for hydration level and thirst)^1,26^. Second, we did not use overt punishments to shape behavior, e.g., air puffs, white noise^11,17,21^. This simplifies the construction of the task and apparatus, and does not introduce interpretational complications of both appetitive and aversive learning in task acquisition and performance. False alarms stayed at low levels in experts, with no need for aversive shaping. Third, by starting operant conditioning for visual stimuli in the most sensitive visual region (binocular space), then gradually introducing increasingly difficult aspects of the task (stimuli in peripheral space, at varying contrast levels, then switching locations), this ensures performance metrics at each training stage can be interpreted according to controlled introduction of new task variables, apart from concurrent learning of the basic stimulus-reward relationship. Fourth, mice learn and perform the task in a semi-enclosed plastic tube, where no locomotion is possible. This is important because locomotion introduces widespread neural modulation across the brain, increases eye movements, impacts perceptual performance^40,41^, and can dramatically alter the strength of visual responses, all potentially complicating the interpretation of attention effects^18,42–44^. In this way, our task design aligns more closely with head-fixed, stationary conditions in monkeys^9,45^, facilitating comparisons of attentional modulation across species.

Our training protocol also carries several inherent limitations resulting from task design choices. A first limitation is that visual stimuli do not appear in symmetric positions in each visual hemifield across blocks of trials^11^, as typically positioned in primate studies^34^. Our choice was for several reasons: 1) to exploit the greater visual sensitivity of binocular visual space during the initial operant conditioning phase; 2) so that we could study the differences in perception (and its improvement by attention) in these two functionally distinct regions of visual space; 3) so that during recordings, stimulus-evoked neural activity in monocular blocks is initially localized to one cerebral hemisphere. Much evidence shows that the binocular visual field in mice is specialized and behaviorally privileged, even in the absence of a fovea^21,22,31–33^. Our task provides a standardized and rigorous way to examine the neural basis of these visual field differences. A second limitation is that our task can only examine the progression of attentional effects across successive trials following the first appearance of the visual stimulus in the unexpected location (block switch). Since the duration of each intertrial interval is randomized, we do not have explicit events (such as an orienting cue) that align the onset of attention in the exact same way on every single trial. Multiple behavioral readouts in our task indicate that mice took several trials to shift their attention, likely accumulating evidence of stimulus location and successive reward outcomes. This slow accumulation across trials (with randomized intervals spanning seconds to tens of seconds) constrains temporal examination of the onset and duration of attentional effects. Modifications to our task with visual cueing of the relevant spatial location could be implemented to study temporally-aligned attentional effects^11^. A third limitation is one common to all detection tasks: there is difficulty in interpreting failed detection trials, relative to two choice tasks^19,46–48^. However, our use of blank stimulus trials with the exact same temporal structure as true stimulus trials shows that false alarms did not increase across successive trials in the block, while hit rates, reaction times, and contrast thresholds all improved — consistent with attentional improvements associated with greater visual sensitivity^37^.

There are several behavioral factors that could be considered in the future to improve training efficiency. For one, we did not account for variations in learning speed across animals, although these were evident (Fig. 2C). These variations may arise from many factors including strain differences^49,50^(but see Fig. S9), breeding paradigms^51^, animal sex^28^, and environmental factors^52^. All of these could be further analyzed and controlled. Second, mice were trained on a reverse-light cycle and at approximately the same hour of day within subject, but we did not examine if circadian effects explain subject-to-subject learning variability and performance^53,54^, a topic for future consideration. Third, trainer sex^55^ and trainer experience can also impact training variability, although both novice and experienced trainers of either sex trained mice to expert-level performance using the protocol (Fig. S5).

Attention is a fundamental cognitive process that influences our vision and perception; our mechanistic understanding of this has come from decades of foundational studies in primates. Our protocol provides an efficient and reproducible way to study spatial attention behavior in head-fixed mice, permitting future use of cell-type specific perturbations, advanced optical and electrical recordings, and use of transgenic mouse models of neurological disorders, many of which show deficits and impairments of attention. By pairing these techniques with rigorous and reproducible procedures for training behavioral tasks, mice can provide detailed insights into the neural basis of attention, and more clearly delineate conserved versus ethologically distinct mechanisms of attention in mammalian sensory systems.

## Resource availability

### Lead contact

Further information and requests for resources should be directed to and will be fulfilled by the lead contact, Bilal Haider.

### Materials availability

This study did not generate new, unique reagents.

### Data and code availability

All data structures and code that generated each figure will be publicly available at DOI 10.6084/m9.figshare.30375733 upon publication and linked from the corresponding author’s institutional webpage upon publication.

## Acknowledgments

We thank members of the Haider lab for feedback. This work was supported by the Alfred P. Sloan Foundation’s Minority Ph.D. (MPHD) Program Fellowship (to J.D.R.), the National Eye Institute (F31EY033691 to K.P.) and the National Institute of Neurological Disorders and Stroke (T32NS096050 to K.P. and R01NS109978 and RF1NS132288 to B.H.), the Simons Foundation (grant no. SFARI 600343, B.H.), and the National Science Foundation Graduate Research Fellowship Program (DGE-2141064 to K.C.J).

## Contributions

Conceptualization: K.P., B.H., Data Curation: K.P., N.A., Formal Analysis: K.P., N.A., Behavior Training: K.P., N.A., J.D., J.Z., K.C.J., M.L-E., E.K., L.L., A.J.O., K.W., J.A., A.D.L., Y.Y., Methodology: K.P., B.H., Supervision: B.H., Validation: K.P., N.A., Visualization: K.P., B.H., Writing: K.P., B.H., Funding Acquisition: K.P., J.D.R, K.C.J., B.H.

## Methods

### Experimental model

All experiments were approved by the Institutional Animal Care and Use Committee at Georgia Institute of Technology and adhered to NIH guidelines. Male (N=30) and female (N=6) mice (8–12 weeks old, Jackson Labs) were used for all studies. Mice were bred in-house and housed in groups of 2–4 in ventilated cages under a 12-hour reverse light/dark cycle with temperature and humidity controlled (21–23°C, 40 – 60%) prior to being selected for experiments. Mice were randomly chosen for experiments, as long as they had no history of poor health, were the appropriate weight for their age, and met genetic requirements (i.e., were the correct genotype). When mice were selected for training, they were kept single housed as previously described^20,33,56^. Enrichment (e.g., nesting materials) was provided in all cages. Food and water were provided *ad libitum* before training; thereafter, water was removed from the cage and hydration followed the protocol as described in our prior studies^20,33,56^ and below.

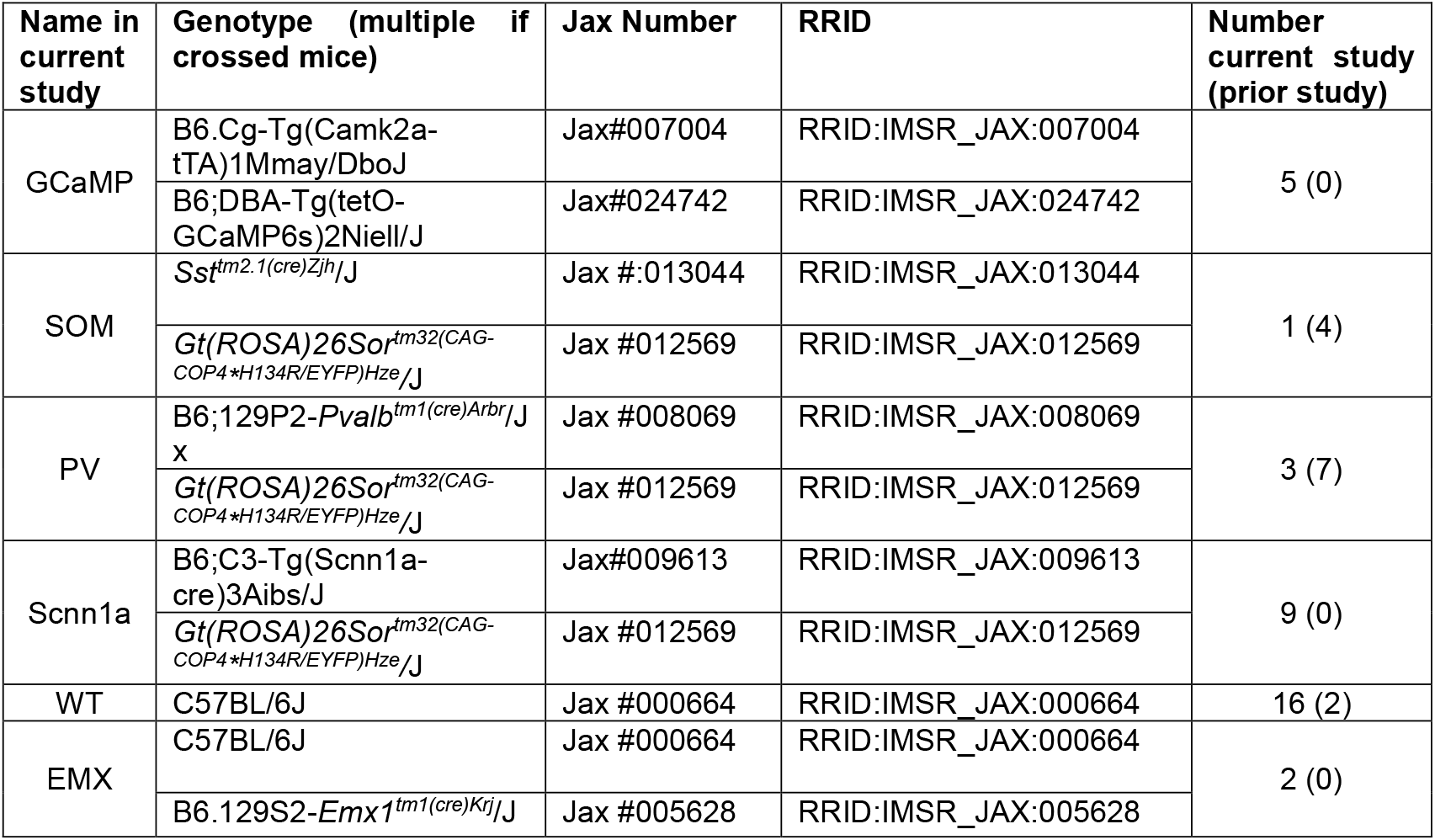

### Headplate implantation

Mice were anesthetized with isoflurane (3% induction, 1-2% maintenance) and implanted with a custom stainless steel head plate with integrated recording chamber as described previously^20,56^. The head plate was attached to the skull using a layer of adhesive (VetBond) before being fully secured to the cranium using dental cement (Metabond). The recording chamber was sealed with an elastomer (KwikCast) to protect the skull. Following implantation, analgesics were administered immediately upon recovery from anesthesia (sustained-release buprenorphine, 1 mg/kg) and mice were given a minimum of three days to fully recover in their home cage.

### Handling and habituation

Mice underwent gentle handling and habituation for at least three days before training. On day one, experimenters handled the mice in their home cages for 15–30 minutes. On day two, mice were handled for 30–45 minutes, and introduced to head-fixation in the home cage, but not placed on the training rig. By day three, the mice were head-fixed and placed on the training rig for 15 minutes, without any stimuli or the lick detector present. Each mouse was trained at the same time every day.

### Water restriction and weight monitoring

The pre-restriction weight of each mouse was recorded as its baseline. Mice were single-housed and water-restricted for 24 hours before the first behavioral task training day, and received an *ad libitum* mix of standard and high-fat diets to maintain weight (50/50 mix of Purina Lab Diet, Laboratory Rodent Diet 5001; Purina Lab Diet, Mouse Diet 5015). Weights were recorded both before and after training, along with the amount of water received during the task. Each correct trial provided 2.5–3.0 µL of water. The minimum amount of water given per day was calculated from the age-and strain-matched minimum expected weight from Jackson labs (typically 1.0mL / day for a 25g mouse). If the daily minimum water amount was not met during the task, mice were supplemented with Hydrogel, administered 60–120 minutes post-training. Hydrogel was administered daily, including weekends. If a mouse dropped below 75% of the minimum weight, as per prior studies ^57^ they were given the day off with full supplemental water and food (e.g., sunflower seeds, high-fat diet). If weight remained below the minimum for more than three days, the mouse was given *ad libitum* water and removed from training (1 of 39 mice in the current study).

### Behavior apparatus

The mouse was head-fixed and rested comfortably on its hindlimbs in a semi-enclosed tube with forelimbs resting on a bar. The mouse was placed in the center of the behavior training apparatus on a custom-built platform (Fig. S7). Visual stimuli were displayed using Matlab 2019b and Psychtoolbox version 3.0.19^58,59^. Two computer screens (gamma-corrected LCD displays; Dell Ultrasharp U2417H or U2419H; 60 Hz refresh rate; peak luminance of 250 cd * m^-2^) were placed at 90° angles from one another. The two displays were positioned such that stimuli at 0° and 70° in azimuthal space have similar viewing angles relative to the central axis of the mouse’s eye. Viewing distance from the mouse to each viewing screen was 21 cm. The training chamber was lined with sound-dampening material to minimize outside noise.

Stimuli presented during behavioral training were controlled using a customized version of Rigbox^60^, which logged stimulus presentation, licking events, reward delivery, and eye tracking through a National Instruments data acquisition (DAQ) board. These signals were acquired and logged at 20kHz. All task events (e.g., licks, reward delivery), stimulus parameters (e.g., stimulus duration, contrast, size, position) and behavioral responses (e.g., trial reaction times, trial behavioral outcome) were acquired for every behavioral training session, and timestamped and stored offline for further analysis.

A custom-built lick detector ^61^ was connected to a Tektronix oscilloscope (TBS1064) and licking events were observed in real time by experimenters. The lick detector stays at a fixed position for the duration of a training session. A water delivery spout was placed at the end of the lick detector, within reach of the animal’s tongue. When animals licked (either in response to the detection of a stimulus, or random licking) the tongue movement is picked up by a break in an infrared (IR) beam inside of the lick detector. The breaking of the IR beam during stimulus presentation triggered the release of a solenoid pinch valve that allowed for water to be delivered from the spout inside of the lick detector.

### Eye camera

Face and eye movements were monitored by illuminating the animal’s face and eye with infrared light (Mightex, SLS-02008-A). The right eye was monitored with a camera (Imaging source DMK 21Bu04.H) fitted with a zoom lens (Navitar 7000) and a long-pass filter (Mightex, 092/52×0.75). The camera was placed ~22 cm from the animal’s right eye. Videos were obtained at 30 Hz. These videos were monitored by the experimenter through the entire duration of training. Saved videos were analyzed for changes in pupil size and position using DeepLabCut^62^.

### Task design

Throughout phases 1–4 (described below), mice were required to withhold licking for 0.5–15 s before stimulus onset. This mandatory quiescence period duration was randomized for each trial (exponential distribution). If the mouse licked during this period, the trial restarted with the same duration of enforced quiescence. Following stimulus onset, a lick during the response window (1s long) provided a water reward (hit trial) and turned off the stimulus. Failing to respond resulted in no reward (miss trial) and the next trial began. The stimulus is removed from the screen once the animal has responded, or after 1s (miss trials). A 0% contrast stimulus (‘blank’) was displayed on 20% of randomly selected trials. Licking during a ‘blank’ trial was recorded as a false alarm, and successfully withholding during a ‘blank’ trial was considered a correct rejection. Even though there was no punishment for licking during these trials, mice continually maintained relatively low false alarm rates during training (Fig. 3C, 4G). We define a trial as one single presentation of a stimulus (any contrast, including 0%); a block is defined as the collection of consecutive trials at a single stimulus position; a session is defined as the collection of blocks presented to the animal without any breaks (typically 100-200 trials total), with animals performing for 2-5 sessions per day. Within each session there are several block switches (3-7 per session) and each block consists of the presentation of the stimulus in one location for a given number of trials. Typically, each block consisted of 10 binocular trials and 20-30 monocular trials; trial numbers were unbalanced because animals tended to prioritize performing mostly binocular trials when both were equally available. A training day consisted of all trials across sessions combined. Animals typically performed 2 – 5 sessions per day, with 350-450 trials per day as experts in the task (369 ± 78, mean ± SD, N=36).

### Task phases

#### Phase 0: Operant conditioning

Mice first learned to associate visual stimuli with water rewards through operant conditioning. High contrast (≥65%) Gabor gratings were presented in the center of the binocular region (0° azimuth). Mice were automatically rewarded with 2.5 – 3.0 µL water 0.7 seconds after stimulus onset, regardless of licking behavior. Reaction times were recorded, and once mice demonstrated a hit rate of ≥ 75% and a visual stimulus-triggered reaction time of ≤ 0.7 seconds (i.e., prior to reward onset) for at least 100-150 trials, automatic water delivery was discontinued, and reward was delivered only upon the first lick during the visual stimulus (operant conditioning). The response window typically started at 2s duration in Phase 0. With maintained hit rates and well-aligned visual reaction times (Fig. S2), the response window was shortened to 1s (to incentivize fast reaction times) and mice progressed to Phase 1.

#### Phase 1: Eccentricity shifting

To train mice to detect stimuli in multiple spatial locations, the location of the Gabor grating was gradually shifted in horizontal (azimuth) space by 10-20° increments starting from 0° (binocular region) and ending at ~70° (monocular region). Mice were presented with a block of 50-100 consecutive trials at the same location. Once mice achieved ≥ 75% correct maintaining a d prime of > 1, the stimulus was shifted by another 10-20°. Mice typically displayed reaction times of 0.4 - 0.6 seconds. Once they maintained accuracy of ≥ 75% correct with a d prime > 1 for a stimulus positioned at 70° for at least 1 training day (312 ± 70 trials performed per day in Phase 1; mean ± SD), they progressed to Phase 2.

#### Phase 2: Contrast shaping

In this phase, the stimulus remained positioned at ~70°, but the contrast was varied trial-by-trial. We defined a set of logarithmically spaced contrasts (not including 0%), and four of these appeared during a given block. The range of contrasts progressively shifted to lower values according to performance, but drew from the set of defined reference contrasts. For example, contrast ranges start at [18, 33, 65, 85%] then decrease progressively from [10, 18, 33, 65%] to [5, 10, 18, 33%] to [2, 5, 10, 18%]. A given contrast range was presented in a block of at least 100 consecutive trials, and psychometric performance (hit rate across contrasts) was assessed after every session. The contrast ranges decreased in order to ensure psychometric performance spanned the full dynamic range of performance (from chance level hit rate at the lowest contrast to >75% hit rate at high contrast), with responses to mid-range contrasts falling between 25 - 75% accuracy. Once mice exhibited stable psychometric performance in the monocular region for 1-2 days (375 ± 108 trials performed per day in Phase 2; mean ± SD), they progressed to Phase 3.

#### Phase 3: Stimulus location switching

Stimuli now alternated between monocular (~70°) and binocular (0°) positions in a blocked trial design to probe the behavioral effects of spatial attention. Blocks of 10 – 30 trials were presented consecutively at one location before switching to the other location without any warning or cueing. Performance metrics (reaction time, accuracy, psychometric performance) were analyzed within and across blocks to evaluate how detection speed and accuracy responded to changes in spatial context. By analyzing behavioral performance (hit rate, d’, reaction time) aligned to the block switches, we could assess if successive trials in the block showed improvements consistent with effects of spatial attention. Since sensitivity and performance were much better in binocular versus monocular visual space (Fig. S3A, B), it was important to make sure the animal did not adopt a strategy of simply ignoring the monocular blocks, and waiting to only perform in the binocular blocks. This was achieved by making the monocular blocks slightly longer (more rewarded trials) than the binocular blocks. The ratio of trials in [binocular: monocular] blocks typically started at [10:20], and was adjusted if needed per mouse based on psychometric performance in each location (average block sizes across N = 36 trained mice; binocular: 12 ± 2 trials; monocular: 22 ± 6 trials, mean ± SD). The most common adjustment was increasing the monocular block length if there was evidence of diminishing performance at that location. Animals were required to maintain psychometric performance with the highest contrast resulting in a d’ value > 1 in both spatial locations to be considered experts. The full behavioral effects of spatial attention could be measured in Phase 3. See Supplementary Video 1 for an example of a mouse performing the task during Phase 3. For animals that were trained for simultaneous neural recordings during behavior, we trained them additionally in Phase 4.

#### Phase 4 (optional): Introduce task-irrelevant bars

We continued training exactly as described in Phase 3, but introduced low-contrast (±5%) task-irrelevant black and white vertical bars (10° wide, full screen height) presented one at a time (duration 0.1s or 0.4s) in random locations throughout the visual hemifield (spanning 160°). These bars were only used to measure neural receptive field responses during recordings, and to probe the influence of spatial attention on those responses. Our previous study showed that these low contrast bars did not interfere with detection of the higher contrast gratings^20^. Bars appeared during the entire behavioral session, during both inter-stimulus intervals and during stimulus events. Initially, bars are introduced at very low contrast (1 - 2% against gray background). If behavioral performance after the introduction of bars remained stable (d’ at highest contrast in both locations ≥ 1, psychometric performance in both locations), then bar contrasts slowly increased (in 1 - 2% increments) over training days until reaching the final 5% contrast value. See Supplementary Video 2 for examples of mice during Phase 4. Phase 4 added an additional 7 ± 9 days of training, on average.

### Subject inclusion/exclusion

A total of 39 mice were selected to be trained in the standardized protocol. Of these mice, 3/39 (8%) were excluded due to: headplate issues (1/39), falling below minimum weight during training (1/39), or failing to reach performance criteria to move on to the next phase (Phase 1 training, failed to reach > 75% hit rate for high contrast monocular trials, 1/39). The remaining 36 mice (89%) satisfied inclusion criteria and completed the full task (Phase 0-3). Of these, 34/36 also completed the optional Phase 4. For Figure 4 (Contrast shaping), two mice were excluded because they only required contrast shaping for a single day, preventing early versus late comparisons across days.

### Experimenter observations and interventions

A detailed description of experimenter observations, interventions, and troubleshooting is described in Supplementary Information.

### Task parameters

Experimenters adjusted the following task parameters according to the protocol: stimulus contrast, location, and the window of time for animals to withhold licking prior to stimulus onset (‘quiescence window’). Quiescence windows were drawn from an exponential distribution (flat “hazard” function), where experimenters defined the minimum, maximum, and median for the distribution. Each trial’s pre-stimulus quiescence window was drawn from this distribution. Longer quiescence windows were used to minimize false alarms (typical range: minimum 5s, maximum 15s, median 10s), while short windows were used when animals achieve expert-level performance in the task (typical range: minimum 1s, maximum 8s, median 5s). Between each session, these quiescence window parameters were randomly adjusted by 1-2 s to prevent animals from adopting a timing strategy. The average quiescence parameters for mice at the expert stage (during neural recordings) were: [minimum: 1.0s; maximum: 6.8s median: 4.5s].

### Visual Stimuli: Gabor Gratings

The task-relevant rewarded stimuli were Gabor gratings with spatial frequency (0.05–0.1 cycles/degree) and size (σ = 10°) fixed across trials, while grating phase was randomized trial-to-trial (to mitigate adaptation). Stimuli were displayed against a 50% gray background.

### Visual Stimuli: Low Contrast Task-Irrelevant Bar Stimuli

During the optional phase 4 of training, low-contrast (2-5%) black or white task-irrelevant vertical bars (10° wide, full screen height) were briefly presented (0.1 s duration, 0.3 s inter-stimulus interval in some cases; 0.4 s duration with no interval in other cases), one at a time, in a given position randomly from −45 to 117° (spanning 160° of the visual hemifield). These vertical bars are the same as used in our previous studies for mapping spatiotemporal receptive fields in mouse primary visual cortex both during and outside of a behavioral task^20,33,56,63^.

### Performance metrics and behavioral analysis

Quantitative metrics used to assess task performance included hit rate, miss rate, false alarm rate, correct rejection rate, reaction time (RT), and psychometric curves to evaluate sensory thresholds. As in our prior studies^20,56^, we used signal detection theory to calculate sensitivity (d prime; d’). Behavioral analysis was automatically generated at the end of every session to track learning and inform experimenter decision making through training stages (Fig. S8). At the end of Phase 3, analysis was aligned to the first trial of block starts in the two stimulus locations (e.g. Fig. 5) to assess effects of attentional shifts with location switching.

#### Quantification of attention effects

Behavioral evidence of attentional improvements included decreased reaction time, and/or increased accuracy and/or d’ on the later trials in the block following the stimulus location switch. To quantify this across the population, we used a linear mixed-effects model (fitlme, Matlab 2024b) with trial number as a fixed effect and random intercepts for mouse and day, allowing us to assess attention-related changes both within and across animals (Fig. 5).

To assess attention effects within individual training days during phases 3 and 4, we compared performance on each day between early trials (1-2) and late trials (6-7) for each mouse. For each day, four metrics were analyzed: reaction time (RT) and detection sensitivity (d’) for both monocular and binocular stimuli. To determine whether the observed effects exceeded what would be expected by chance, a permutation test was performed for every training day and metric across mice. Trial order was randomly shuffled 1000 times, and for each shuffle the difference between late and early trials was computed. The 95% confidence interval of this permutation-derived null distribution defined the range expected under the null hypothesis of ‘no attention effects.’ A day was labeled as a significant attention day if the observed late-early difference for any of the four metrics fell outside of this 95% confidence interval in the expected direction (e.g., decreases in the RT, increases in the d’).

In addition, to control for a potential confound of subject-to-subject differences in absolute reaction time range or d’ range driving the effects, we used a modulation index (MI) within subjects to calculate the relative change in reaction time and d prime between late (trials 6-7) and early (trials 1-2) trials within each day (Fig S1), defined as:

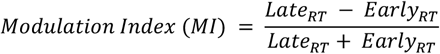

Because d prime values can be negative, the equation to calculate the MI for d prime changes across trials within a day was:

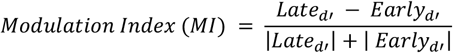

#### Perceptual contrast sensitivity: psychometric curve fitting

The hit rate (proportion of correct trials) was measured across contrasts. A Boltzmann’s sigmoidal equation was used to fit the data points to generate psychometric contrast response functions:

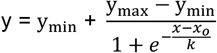

y_min_: minimum of the output variable (false alarm rate)

y_max_: maximum of the output variable (hit rate at highest contrast)

*x:* contrast values

*x*_*o*_: midpoint of the sigmoid curve (used to extract contrast threshold, C50)

*k:* steepness parameter

The data was fit using the MATLAB function ‘fit.’

#### Perceptual contrast sensitivity: attentional effects following block switch

Psychometric sensitivity was analyzed across successive trials following a block switch. The label ‘monocular’ indicates all the trials following a switch from a binocular to monocular grating location. Likewise, the label ‘binocular’ indicates all the trials following a switch from a monocular to binocular grating location. In either case, behavioral data in phases 3 or 4 of training were aligned to the block switch, and hit trials aggregated across all block switches in all subjects. Since contrast levels for each trial were randomized (e.g., the first trial might display a stimulus at 10% contrast in one block and 18% contrast in another), all stimuli across all of the blocks can be used to construct a contrast responses curve from *all* of the first trials after the block switch. These aggregate psychometric curves were constructed for each successive trial and illustrate how the psychometric contrast sensitivity across all contrasts changes following the block switch (Figure 6A, C). Contrasts were converted to proportions (0-1) prior to fitting and slope calculation. Contrast thresholds and slopes were derived from the psychometric curves, following a similar approach outlined in^34^.

Psychometric curves were fit separately for monocular and binocular conditions using a four-parameter Boltzmann function (see above). Data were organized by animal, day, and trial number following a block switch. For some days, curve fits could not be computed for certain trial–contrast combinations due to a lack of trials (274 of 759 total days); these were excluded from further analysis. Only trials from days where sensitivity (d’) increased as a function of trial number within the block (monocular or binocular) were included, resulting in 234 of 485 total days. From each successfully computed fit, contrast threshold (C50) and slope parameters were extracted. Linear mixed-effects models (fitlme, Matlab 2024b) were used to assess changes in these parameters across trials after the block switch, including random intercepts for animal, and day nested within animal. Group averages (mean) and standard errors (SEM) were computed for visualization across successive trials after the block switch (Fig. 6).

---

**Figure S1.**
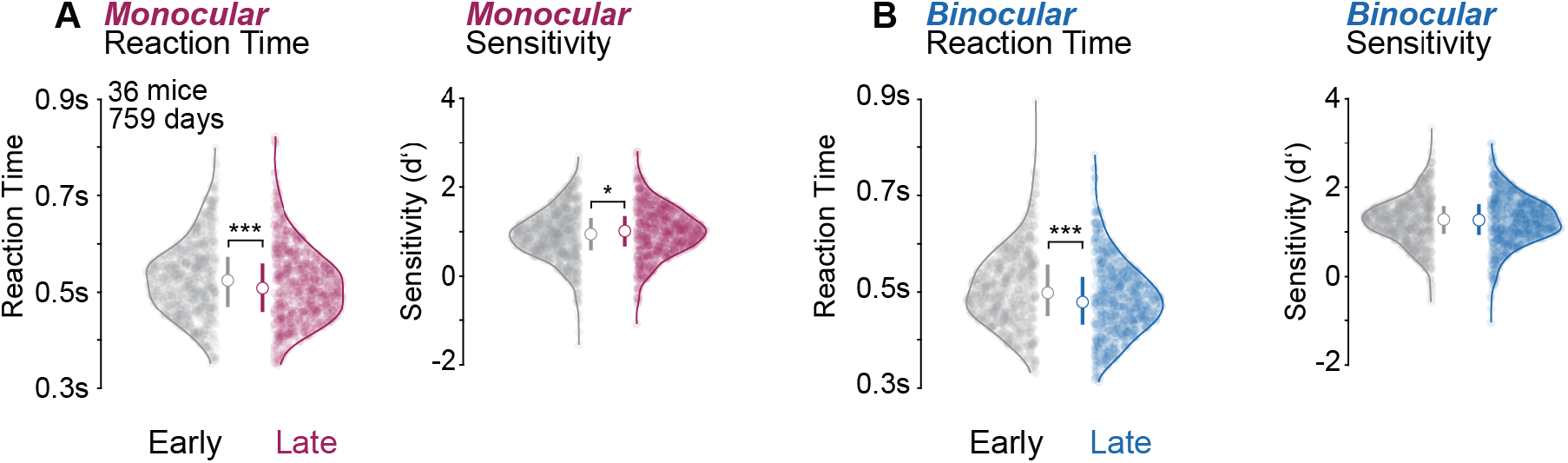
Protocol trained mice show improved detection performance following block switches. **A**. Mice trained in the optimized protocol (N=36 mice, 759 total days) exhibit faster reaction times (*left*, Early: 0.52 ± 0.05s; Late; 0.50 ± 0.05s; median ± IQR/2 throughout figure; *p =* 0.0006, sign rank test) and improved sensitivity (right; Early: d’ = 0.95 ± 0.35; Late: d’ = 1.02 ± 0.34; *p =* 0.021, sign rank test) for stimuli in monocular space during later trials (trials 6-7) relative to early trials (trials 1-2) after a block switch. These findings replicate effects reported in Speed et al., 2020. Additional analysis using a modulation index (MI) within subjects that controls for individual differences in the range of reaction times or sensitivity (see Methods) also finds sign9ificant effects of late versus early reaction times and d prime values per day (reaction time MI: *p =* 6.41e-11; d prime MI: *p =* 8.38e-05, sign rank test). **B**. Mice trained in the optimized protocol (N=36 mice, 759 total days) also exhibit faster reaction times for stimuli in binocular space during later trials relative to early trials after a block switch (left; Early: 0.50 ± 0.54s; Late: 0.48 ± 0.05s; *p =* 7.63e-8, sign rank test). Binocular sensitivity did not improve in later trials relative to early trials (right; Early: d’ = 1.31 ± 0.31; Late: d’ = 1.3 ± 0.33; *p =* 0.904, sign rank test). Additional analysis using a MI of late versus early reaction times and d prime values per day reveals similar results (reaction time MI: *p =* 1.15e-24; d prime MI: *p =* 0.955, sign rank test).

**Figure S2.**
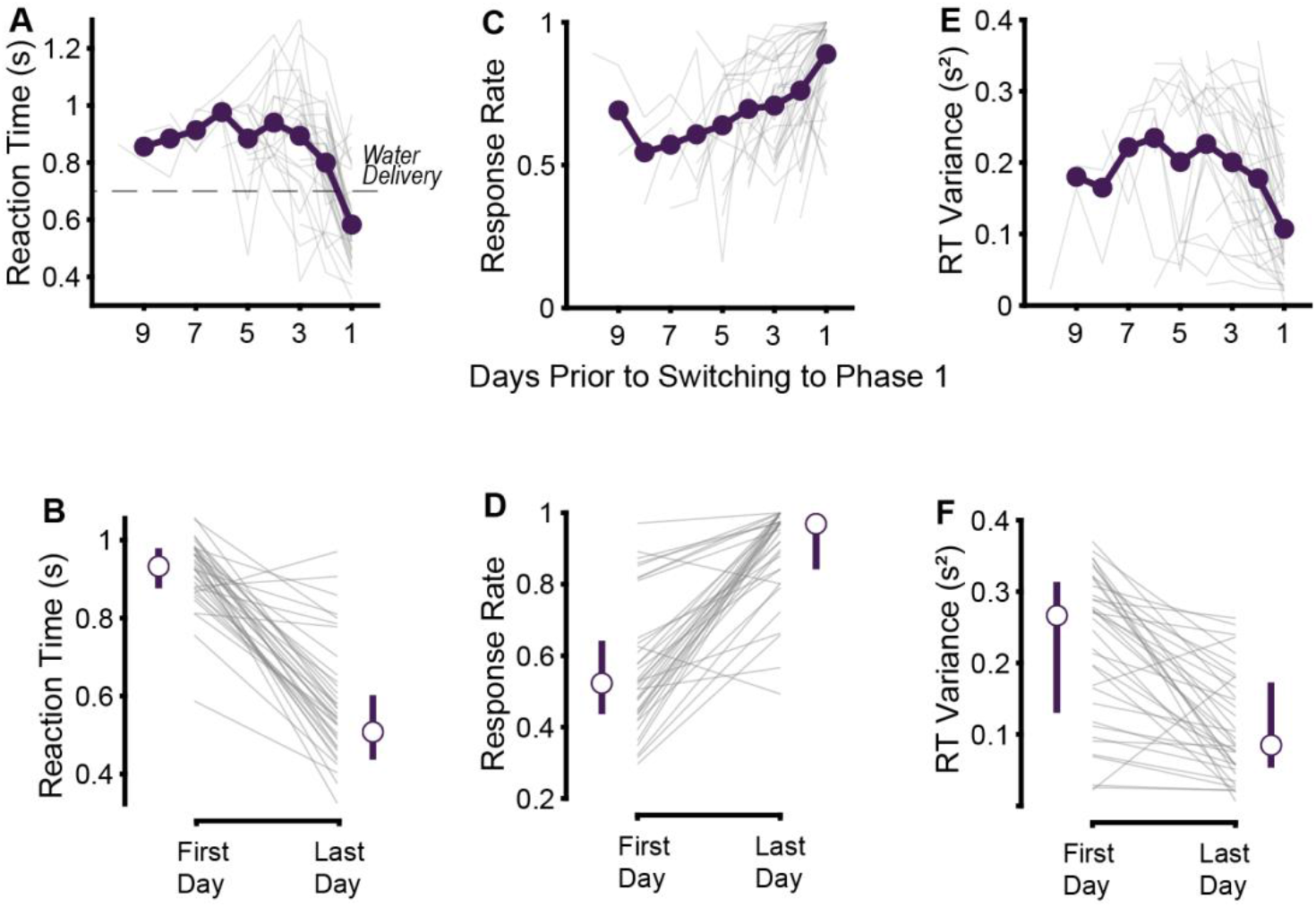
Operant conditioning establishes robust stimulus-reward association. **A**. Reaction times become quicker as animals progress through Phase 0 (3-9 days). Prior to the transition to Phase 1 of training, mice exhibit reaction times that occur before the automatic reward delivery (< 0.7 s, dashed line). **B**. On the first day of training, reaction times are significantly slower (0.93 ± 0.05 seconds after stimulus onset, median ± IQR/2 throughout figure) than on the last day of training (0.55 ± 0.08 seconds after stimulus onset; *p =* 2.56e-7, sign rank test) in Phase 0. **C**. Response rate increases as animals progress through Phase 0. Animals reach 75% or greater accuracy for detecting stimuli prior to transitioning to Phase 1. **D**. On the first day of training, response rates are significantly lower (52.0 ± 10%) than on the last day of training (97.3 ± 7.5%; *p =* 3.57e-7, sign rank test) in Phase 0. **E**. Reaction time variance decreases as animals progress through Phase 0 and reaction times become more reliable to presentations of binocular stimuli. **F**. On the first day of training, reaction time variance is significantly higher (0.27 ± 0.09 s^2^) than on the last day of training (0.08 ± 0.05 s^2^, *p =* 3.31e-06, sign rank test) in Phase 0.

**Figure S3.**
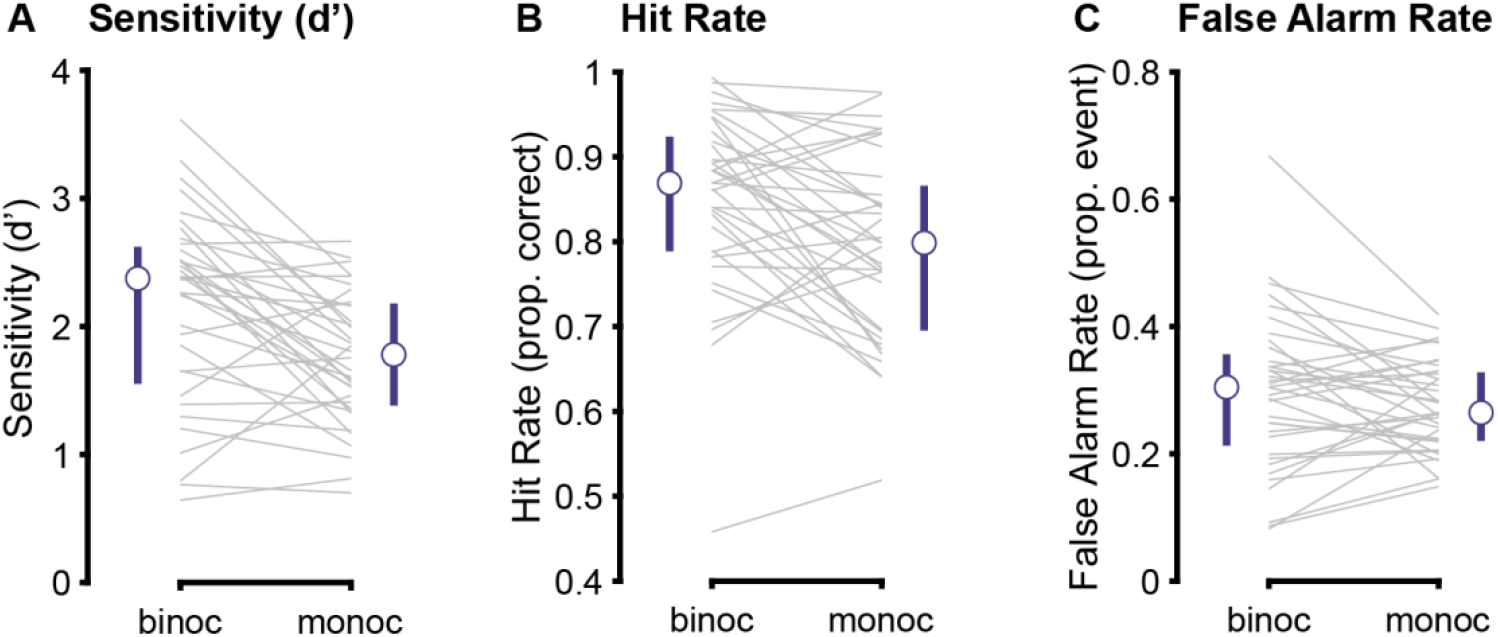
Sensitivity, hit rate, and false alarm rate across individual mice during binocular and monocular stimulus detection sessions during Phase 1 training. **A**. Sensitivity (d prime, d’) to binocular (binoc; ≤ 40° azimuth) stimuli was significantly higher than sensitivity to monocular stimuli (monoc; > 40° azimuth) during Phase 1 training (*p =* 0.003, sign rank test). During Phase 1, stimulus contrast did not differ between stimulus locations. 21/36 mice were presented with 65% and 85% contrasts for both binocular and monocular stimuli; 15/36 were presented with 85% contrast only. **B**. Binocular hit rates significantly higher than monocular hit rates during Phase 1 training (*p =* 0.004, sign rank test). **C**. False alarm rates were not significantly different for binocular versus monocular stimulus detection in Phase 1 (*p =* 0.132, sign rank test).

**Figure S4.**
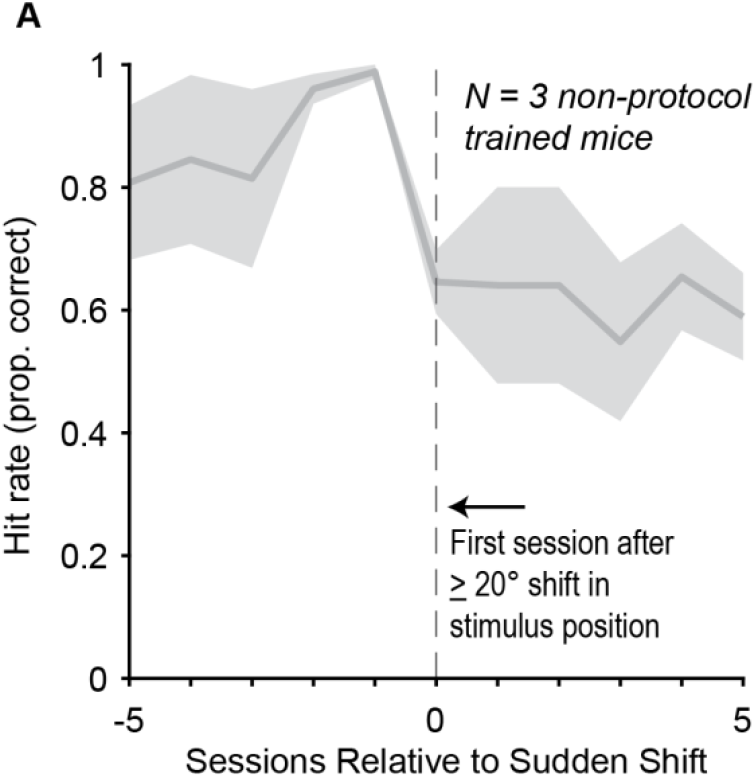
Sudden shifts in stimulus position during Phase 1 training result in poor detection performance. **A**. In N=3 mice not trained in the protocol, a sudden ≥20° shift from binocular to monocular visual space resulted in an average 34% reduction in accuracy. Hit rates for individual mice decreased as follows: Mouse 1, 100% to 60% (shift from 30° to 50°); Mouse 2, 100% to 58% (30° to 60°); Mouse 3, 97% to 75% (40° to 60°).

**Figure S5.**
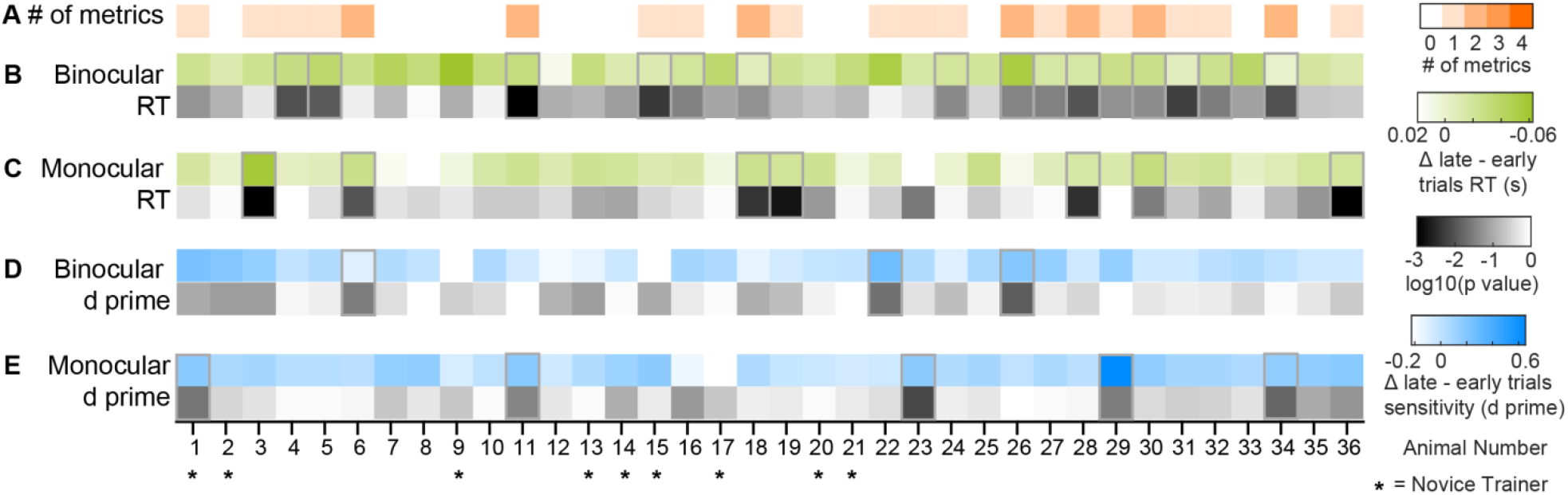
Changes in reaction time and sensitivity on trials after a block switch for all mice. FRigeulraeteSd6t.oPFuipgiul rseiz5e. and position do not change across trials after a block switch. **A**. Number of statistically significant metrics of visual spatial attention (4 metrics defined in B-E), averaged across all Phase 3 and 4 days for each animal (no selection criteria for sessions or days; p < 0.05, permutation test, see Methods). In the average of all Phase 3 and 4 days, 61% of all mice showed at least one significant metric, and 83% of mice showed at least one day with statistically significant effects in Phase 3 or 4. At the single day level, overall 15 ± 10% of all Phase 3 and 4 training days showed statistical significance of at least one metric (mean ± SD across animals), suggesting that the direction of effects on individually non-significant days nonetheless remained consistent and passed significance threshold with cumulative samples (Fig. 5). **B**. Average difference of reaction times (green) between late (6–7) versus early (1–2) trials across the block, averaged for all Phase 3 or 4 training days for each mouse (columns) during binocular blocks. Darker green indicates faster reaction times later in the block. Log p-values (Wilcoxon signed-rank test) in greyscale for each mouse (significant subjects outlined in top row). **C**. Same as B, but for monocular blocks. **D**. Same trials as B, but for early versus late differences in binocular sensitivity (d’). Darker blue indicates a larger difference. Log p-values denote significance (Wilcoxon signed-rank test). **E**. Same trials as C, but for early versus late differences in monocular sensitivity (d’). Stars (*) indicate mice trained by a novice trainer following the protocol for the first time.

**Figure S6.**
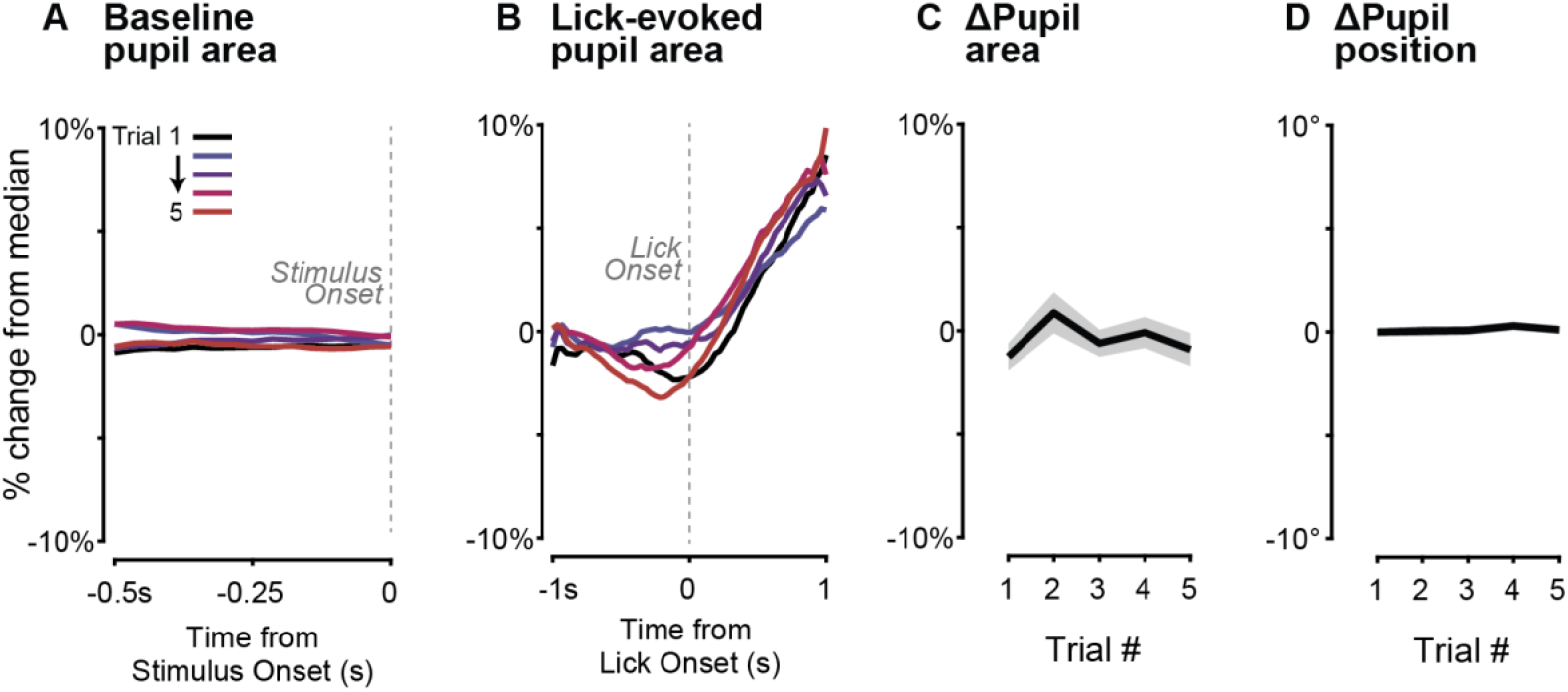
Pupil size and position do not change across trials after a block switch. **A**. Percentage change in baseline pupil area (relative to the median pupil size across the entire session) before stimulus onset, sorted by successive trials following location switch (Trials 1 to 5, colored), for all subjects with high quality pupil videos (n = 25 mice). **B**. For comparison to A, pupil size reliably increases >10% relative to the baseline after lick onset and reward. **C**. Pre-stimulus pupil area (from A) does not change significantly across successive trials after a block switch (RM ANOVA, *p =* 0.164; Trial 1: −1.25 ± 0.77% versus Trial 5: −0.9 ± 0.95%, mean ± SEM; n = 25 mice, 1063 correct detection trials, 137 block switches (both monocular and binocular), and 26 sessions). **D**. In the same trials, pupil position prior to stimulus onset does not change significantly across trials after a block switch (RM ANOVA, *p =* 0.750; Trial 1: −0.01 ± 0.22° versus Trial 5: 0.1 ± 0.23°).

**Figure S7.**
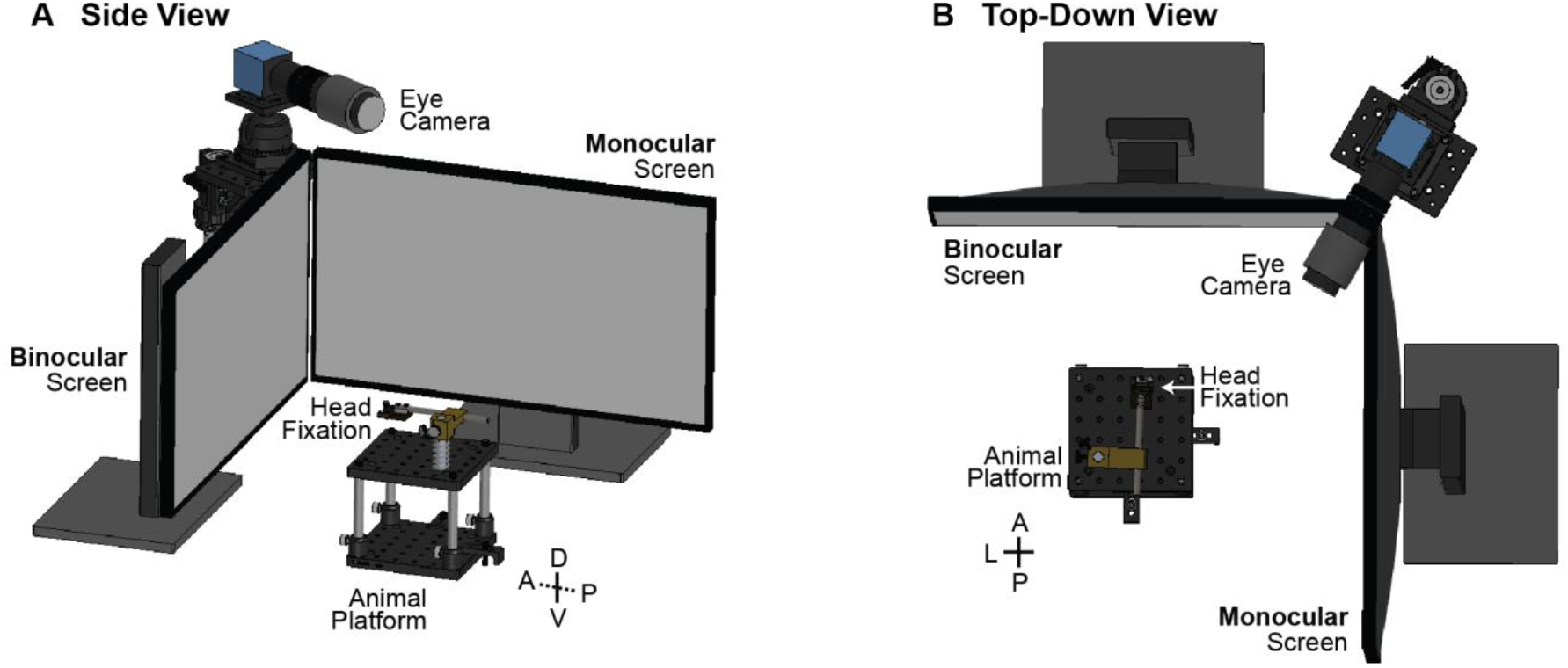
Mouse behavior training setup. **A**. Side view of mouse behavioral training setup. Mice are head fixed on the animal platform facing the binocular screen. The monocular screen is always placed to the right of the animals. The eye camera normally points towards the face and eye during training. **B**. Same as in A, but top-down view of the behavioral training setup.

**Figure S8.**
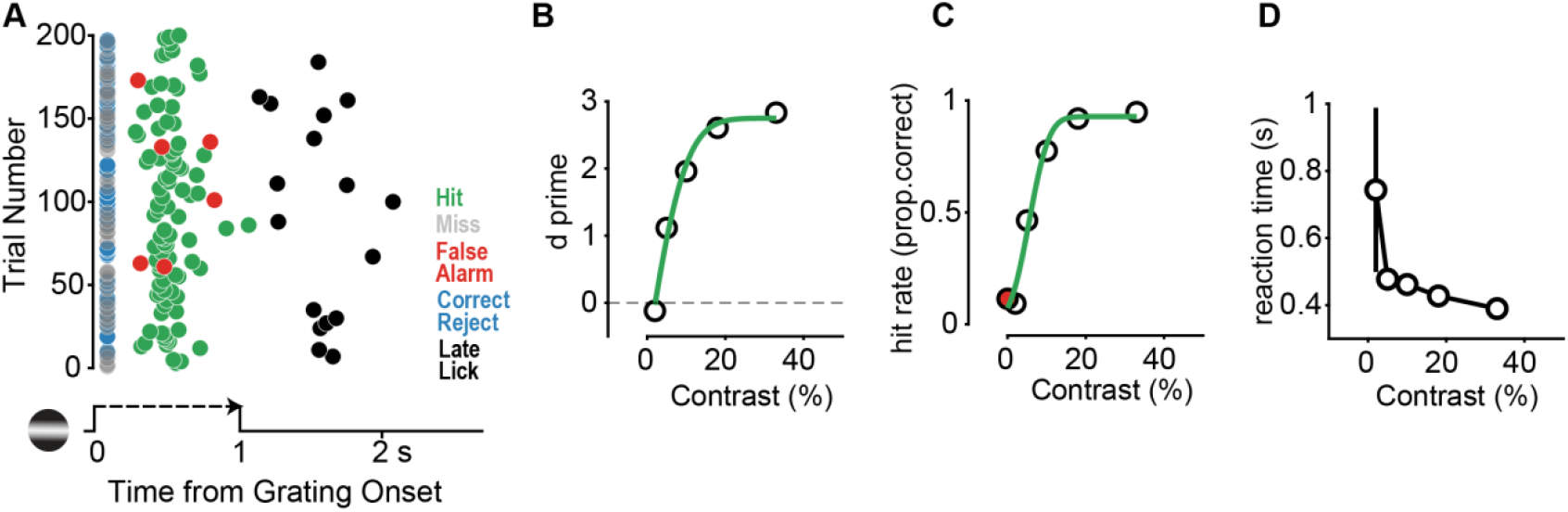
Session-by-session plots viewed by experimenters. **A**. Example performance of a mouse in a single behavior session within a day (monocular performance, Phase 2, Contrast shaping). Reaction time (latency of first lick after stimulus onset, x-axis) for the detection of stimuli on each trial (y-axis). Each trial is labeled with a colored circle depending on the trial outcome: successful detection (hit; green), failure in detection (miss; gray), false alarm (red), correct reject (blue), and a late response (responding within 1 s after stimulus offset, black). Detection failures and correct rejects (both trial types with no lick responses) arbitrarily plotted at 0 sec. **B**. Mean sensitivity (d’) at each stimulus contrast value presented in the session (open circles). Green line indicates Boltzmann fit. **C**. Mean hit rate (proportion of correct responses) at each stimulus contrast value presented in the session (open circles). False alarm rate (responses to 0% contrast stimuli) is indicated by the red circle. **D**. Mean reaction time at each stimulus contrast value presented in the session (open circles). Error bars represent standard deviation.

**Figure S9.**
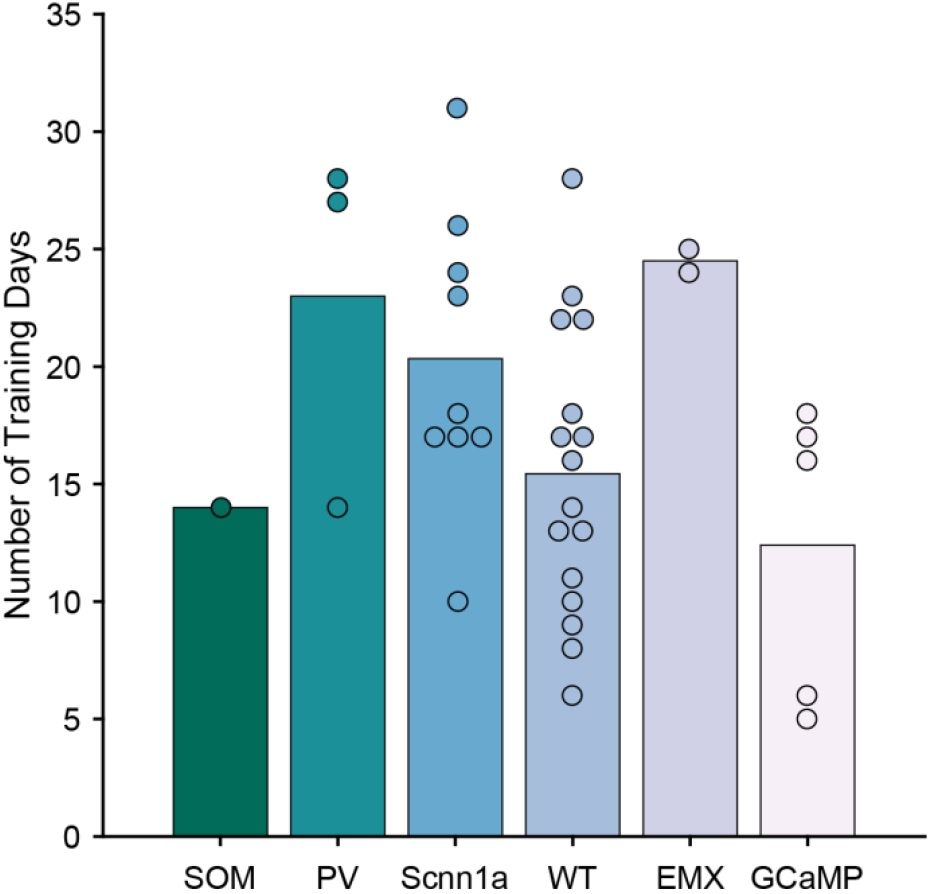
Time to train does not significantly differ across genotypes. **A**. Six genotypes of mice were used during training. Average time to train for each genotype are as follows: SOM (green, N=1, 14 ± 0 days; mean ± SD throughout figure); PV (turquoise, N=3, 23 ± 8 days); Scnn1a (light blue, N=9, 20 ± 6 days); WT (periwinkle, N=16, 16 ± 6 days); EMX (lavender, N=2, 25 ± 1 day); GCaMP (light pink, N=5, 12 ± 6 days). There was no significant difference in total training time (up to Phase 3) across genotypes (*p =* 0.075, Kruskal–Wallis test). Full names and details for each strain are listed under Methods: Experimental Model.

**Supplementary Table 1.** Related to Figure 5, Linear Mixed Effects Model Details.

| Measure | $\beta$ (Estimate ± SE) | t-statistic (df) | p-value | Model |
| --- | --- | --- | --- | --- |
| Binocular d' | 0.0012 ± 0.0026 | 0.46 (5304) | 0.64 | d' ~ TrialNumber + (1\|Animal) + (1\|Animal:Day) |
| Monocular d' | 0.0107 ± 0.0027 | 4.02 (5311) | 6.0e-5 | d' ~ TrialNumber + (1\|Animal) + (1\|Animal:Day) |
| Binocular RT | -0.0044 ± 0.0004 | -11.51 (5269) | 2.85e-30 | RT ~ TrialNumber + (1\|Animal) + (1\|Animal:Day) |
| Monocular RT | -0.0026 ± 0.0004 | -6.67 (5305) | 2.9e-11 | RT ~ TrialNumber + (1\|Animal) + (1\|Animal:Day) |

**Supplementary Table 2.** Related to Figure 6, Linear Mixed Effects Model Details.

| Measure | $\beta$ (Estimate ± SE) | t-statistic (df) | p-value | Model |
| --- | --- | --- | --- | --- |
| Binocular C50 | -0.0056 ± 0.0011 | -4.90 (832) | 1.17e-6 | C50 ~ TrialNumber + (1\|Animal) + (1\|Animal:Day) |
| Monocular C50 | -0.0129 ± 0.0027 | -4.75 (806) | 2.38e-6 | C50 ~ TrialNumber + (1\|Animal) + (1\|Animal:Day) |
| Binocular slope | 0.068 ± 0.0315 | 2.15 (832) | 0.032 | Slope ~ TrialNumber + (1\|Animal) + (1\|Animal:Day) |
| Monocular slope | 0.078 ± 0.055 | 1.41 (806) | 0.16 | Slope ~ TrialNumber + (1\|Animal) + (1\|Animal:Day) |

**Supplementary Table 3.** Troubleshooting.

| Observation | Potential Cause | Solution |
| --- | --- | --- |
| Foam in eye, squeaking | Signs of stress | Adjust the animal to make sure they are comfortable and not being hindered by head fixation or positioning. Check lick detector positioning. If signs of stress persist, immediately remove the animal and place in home cage. |
| Slow reaction times relative to usual behavior | Improper animal positioning or lick detector positioning | Make sure the animal is properly positioned in the rig and ensure the lick detector is not touching the animal's face or whiskers. Mark the lick detector position so that it is in the same exact location across days. Also ensure that the animal is in the exact same position daily by marking the location of the mouse tube and marking set points for head fixation system. |
| High false alarm rates | Lick detector too close | Move lick detector away slightly, as it may be too close and the animal engages with the object in front of them. |
|  | High motivation (early in training day) OR animal had previous success with random licking strategy | Increase quiescence time [min: 5s; max: 15s; median 10s] so animal learns to wait to lick until a stimulus appears. |
| Animal will no longer perform the task after a switch from Phase 0 to Phase 1 | Switch from Phase 0 to Phase 1 occurred too early, OR switch from Phase 0 to Phase 1 | Introduce Phase 1 during the first session of the next training day when motivation for reward is high. If animal still will not perform the task, switch back to Phase 0 parameters |
| | occurred at end of day when animal has low motivation for reward | until animal is reliably detecting the stimulus prior to reward delivery ($< 0.7$ s). |
| Performance crashes at the shift from binocular to monocular locations (e.g., from 40 to 50 degrees) | Increasing difficulty due to more eccentric stimulus presentation | If animal has already consumed $>50\%$ of daily minimum of reward, or performed many trials, then keep the stimulus at the easier location for the duration of the training day. Then the next day when motivation is high, present the stimulus at the slightly more difficult location. Only shift the stimulus by 10 degrees at a time until the animal reaches performance metrics (i.e., $> 75\%$ hit rate) |
| Animal will not detect stimuli in monocular blocks, but will successfully switch to detecting binocular stimuli. | Binocular trials are easier and therefore favored by the animal | 1) Increase the ratio of monocular to binocular trials (e.g., 30M:10B); 2) Decrease binocular contrast range and/or increase the monocular contrast range (do one; assess; then do the other); 3) Give a single session of monocular-only trials ($\sim 50$ stimulus presentations) before re-introducing binocular trials. |

## Supplementary Methods

### Experimenter observations

Experimenters stayed in the same room as the mice for an entire training day. Experimenters trained only one mouse at any given time. During any training session, experimenters were not aware of what contrast or trial type was being shown to the animal on a given trial to minimize interference. Experimenters were able to observe in real-time on an oscilloscope: licking activity; photodiode activity of visual stimulus events; reward triggers. Videos of the face and eye were also recorded to observe pupil, face movements, and assess comfort, engagement, or stress levels of the mouse. After every session, experimenters combined observations of licking activity with the behavioral output (Fig. S8) to adjust task parameters, if necessary.

### Experimenter interventions

Mice typically performed a full behavioral session (50-200 trials) without experimenter intervention. The only real-time interventions experimenters made were to re-adjust the animal if sub-optimally positioned within the recording chamber, or to adjust the lick detector placement if observations indicated this was needed (e.g., the animal pushed it and reaction time latencies suddenly increased). These interventions were always made outside of the animal performing the task (i.e., the experimenter stopped the current stimulus set). These interventions typically occurred earlier in training and were rare later in training as mice gradually became more comfortable in the training environment. A session was stopped if the animal failed to respond for 7 trials in a row (regardless of trial type), or if the animal showed signs of discomfort. During such rest periods (~5 mins), single free rewards were given periodically to assess animal motivation (by examining lick rate evoked by free water reward consumption). Low lick rates indicated low motivation, while high lick rates (~8 Hz) indicated the animal was still motivated by water reward and the session was restarted. If there was still lack of task engagement or continued signs of discomfort, animals were removed from the training apparatus and placed back in their home cage. Mice performed the task until they completely stopped licking (satiety), or reached their full water intake for the day, whichever came first.

## Supplementary Videos

**Supplementary Video 1. Phase 3 Training Video**

**Supplementary Video 2. Phase 4 Training Video**

**Supplementary Video 3. Task Expert Video**

